# Engineering rice Nramp5 modifies cadmium and manganese uptake selectivity using yeast assay system

**DOI:** 10.1101/2024.08.18.605937

**Authors:** Junji Inoue, Takamasa Teramoto, Tomohiko Kazama, Takahiro Nakamura

**Affiliations:** Faculty of Agriculture, Kyushu University, Fukuoka, Japan

**Keywords:** OsNramp5, rice, cadmium, manganese, transporter, selectivity, protein engineering

## Abstract

Cd is a seriously hazardous heavy metal for both plants and humans and international regulations regarding Cd intake have become stricter in recent years. Three-quarters of the Cd intake comes from plant-based foods, half of which comes from cereals. Therefore, it is anticipated that the Cd uptake efficiency of cereals, including rice, a staple crop in Asia, will be reduced. Natural resistance-associated macrophage protein (Nramp) is the principal transporter involved in the uptake and translocation of metal ions in various plants. In rice, OsNramp5 is a transporter of Mn, which is an essential micronutrient for plant growth, and is responsible for Cd uptake. Although several attempts have been made to engineer the metal uptake characteristics of OsNramp5, in many cases, both Cd and Mn uptake efficiencies are impaired. Therefore, in this study, we engineered OsNramp5 to reduce Cd uptake while retaining Mn uptake efficiency for low-Cd rice production. OsNramp5 was engineered using amino acid substitution(s) at the 232^nd^ Ala and 235^th^ Met of OsNramp5, which have been suggested to be key residues for metal uptake efficiency and/or selectivity by structural analyses of bacterial Nramps. The metal uptake efficiency was first analyzed using a yeast model assay system. Several mutants showed less than 8.6% Cd and more than 64.1% Mn uptake efficiency compared to the original OsNramp5. The improved metal uptake characteristics were confirmed by direct measurement of the metal content in the yeast using inductively coupled plasma optical emission spectroscopy. Notably, several mutants reduced Cd uptake efficiency to the background level while retaining more than 64.7% Mn uptake efficiency under conditions mimicking heavily polluted soils in the world. In addition, computational structural modeling suggested requirements for the spatial and chemical properties of the metal transport tunnel and metal-binding site, respectively, for Cd/Mn uptake efficiency.

## 1 Introduction

All organisms require metal ions as micronutrients to maintain their biological activities. Plants absorb biologically useful metals, including Mn^2+^, Fe^2+^, Co^2+^, Ni^2+^, Cu^2+^, and Zn^2+^, from the soil but incidentally incorporate the toxic heavy metals Cd^2+^, Pb^2+^, and Hg^2+^, which pose significant risks to agricultural and human health. Cd is a hazardous heavy metal that can be easily absorbed and accumulated in plant tissues (Gill and Tuteja, 2011). Cd inhibits plant growth by reducing photosynthesis and oxidative stress (Nazir et al., 2022). Moreover, Cd is incorporated into the human body through the food chain and permanently accumulates primarily in the kidneys, with a long biological half-life of 10–35 years (World Health Organization, 2022). High Cd intake can lead to serious health issues, such as kidney dysfunction, cancer, bone fractures, and itai-itai disease (Nogawa and Kido, 1993; Staessen et al., 1999; Tim et al., 2006; Vervaet et al., 2017). Low to moderate Cd exposure is also associated with kidney dysfunction, cancer, and bone density (Järup and Alfvén, 2004; Julin et al., 2012; García-Esquinas et al., 2014; Wallin et al., 2021).

Cd intake has been strictly regulated owing to its hazardous effects. International standards established a permissible maximum intake of Cd at 7 µg/kg/week (1.00 µg/kg/day) in 1988, which was modified to 25 µg/kg/month (0.83 µg/kg/day) in 2010 (World Health Organization, 2011). Europe implemented a stricter standard for intake of 2.5 µg/kg/week (0.36 µg/kg/day) in 2009 (EFSA, 2009). Cd intake is largely caused by seven commodity groups: rice, wheat, roots, tubers, leafy greens, other vegetables, and mollusks (World Health Organization, 2006), six of which are plant-derived. Indeed, it has been reported that Cd intake mostly comes from plant-based foods (74% and 82% in China and Sweden, respectively) and almost half of Cd intake is from cereals: 46% and 48% in China and Sweden, respectively (Julin et al., 2012; Qing et al., 2023). Thus, there is a major challenge in reducing Cd uptake by rice, which is a staple crop for Asian people.

Numerous studies have been conducted to elucidate the molecular mechanisms of Cd uptake in various plants and have found a significant role in natural resistance-associated macrophage proteins (Nramps). Nramp is a principal transporter for the uptake and/or translocation of metal ions (Socha and Guerinot, 2014; Clemens and Ma, 2016). Genomic analysis has shown that plants possess a significantly larger number of *Nramp* genes than mammals and bacteria (Chen et al., 2021). In rice, seven Nramp genes have been found and *Oryza Sativa Nramp5* (*OsNramp5*) was identified to be largely responsible for the uptake of Cd and Mn via roots from the surrounding environment (Ishimaru et al., 2012), as evidenced by *OsNramp5* knockout lines exhibiting impaired Mn/Cd uptake efficiency (Ishikawa et al., 2012; Sasaki et al., 2012).

Various attempts have been made to reduce Cd uptake efficiency by focusing on Nramps. Simple knockdown of *Nramp* genes resulted in a reduction in Mn and Cd uptake and had negative effects on growth in Mn-deficient environments (Cailliatte et al., 2010; Yang et al., 2014; Honma et al., 2017; Gao et al., 2018). This is because Mn is an essential micronutrient for plant growth and development, involving several metabolic pathways, including photosynthesis, and as a cofactor of enzymes. A single amino acid substitution mutant of OsNramp5 Q337K mutant identified through targeting induced local lesions in genomes screening showed a 50% reduction in both Mn and Cd uptake. Notably, a field test of rice containing the Q337K mutation exhibited lower Cd uptake than the wild type and higher growth tolerance to Mn deficiency than knockout lines (Kuramata et al., 2022). Another amino acid substitution mutant, A512T OsNramp5, which was obtained by random mutagenesis, showed a lower Cd uptake efficiency but maintained Mn uptake efficiency in a yeast model assay system, although the evaluation in rice plants has remained (Qu and Nakanishi, 2023). Studies on Nramps involved in Mn/Cd uptake in other plants have suggested that metal uptake selectivity varies depending on the protein species, for example, differences in a few amino acids (Curie et al., 2000; Thomine et al., 2000; Cailliatte et al., 2010).

Structural analyses of bacterial Nramps provide valuable information regarding the molecular basis of metal binding and selectivity (Ehrnstorfer et al., 2014; Bozzi et al., 2016b; Ray et al., 2023). The Nramp family consists of 11 or 12 transmembrane regions (TMs) and the broken helical regions in TM1 and TM6 comprise the substrate-binding sites. Structural studies also revealed critical residues for metal binding (#2, #5, #8, and #11 in Fig. 1) in TM1 and TM6. Several attempts have been made based on structural information and several mutant Nramps with altered metal uptake selectivity (Bozzi et al., 2016a; Li et al., 2018; Lu et al., 2018). However, structure-based engineering of OsNramp5 has not yet been performed.

**Figure 1.**
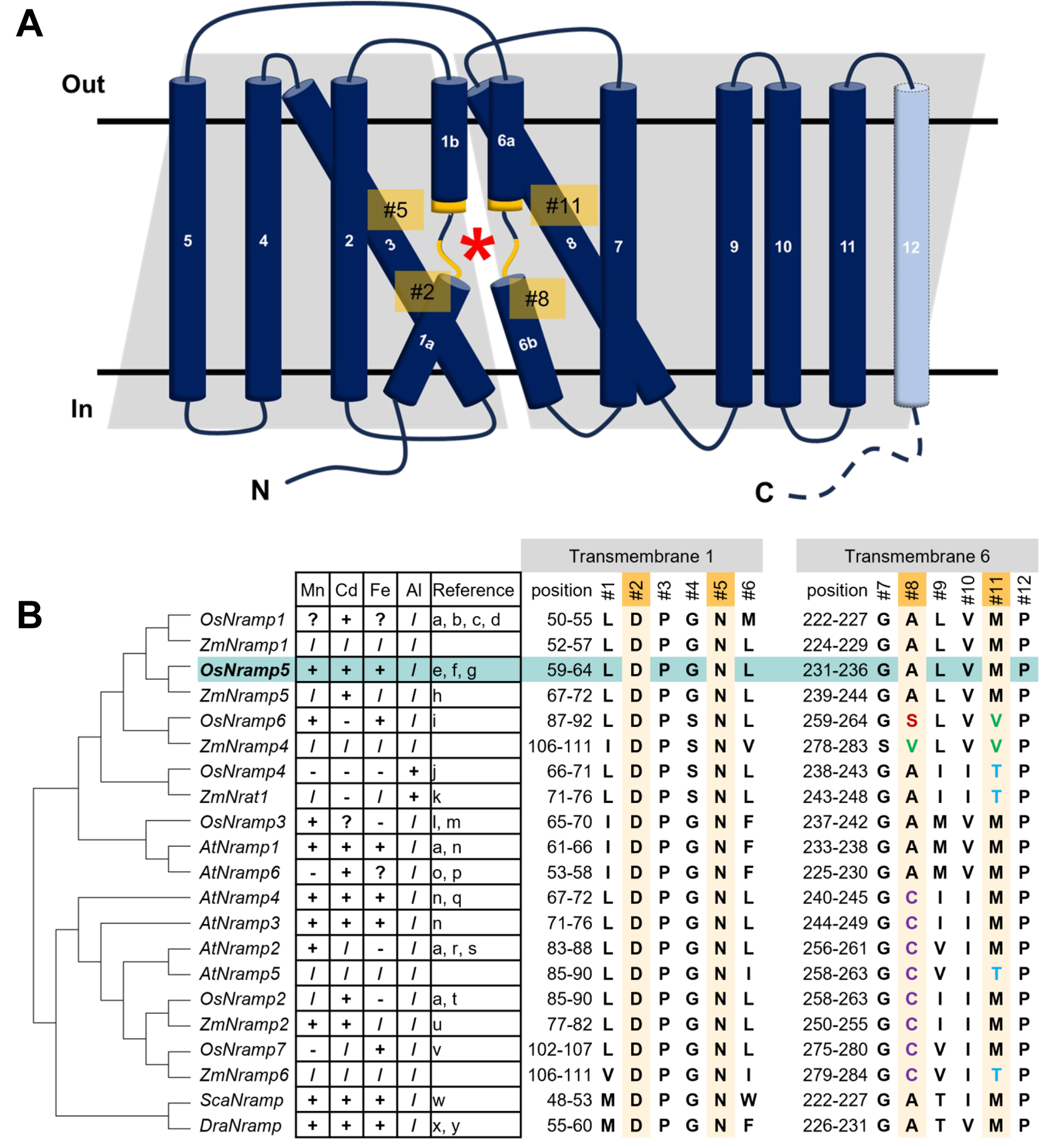
Structure of OsNramp5 and putative amino acids of the metal-binding site involving metal uptake efficiency and selectivity. (A) Topological diagram of OsNramp5 structure. The diagram was depicted based on the domain architecture of UniProt Q8H4H5, with modifications of Figure. 2A in Bozzi and Gaudet, 2021. The OsNramp5 consists of 12 transmembrane regions. The asterisk denotes the location of the metal-binding site composed of TM1 and TM6. The putative key amino acid residues for the metal binding are designated as #2, #5, #8, and #11. (B) Phylogenetic tree and amino acid composition of metal-binding sites of 21 Nramps in various organisms (Os, *Oryza sativa*; Zm, *Zea mays*; At, *Arabidopsis thaliana*; Sca, *Staphylococcus capitis*; Dra, *Deinococcus radiodurans*). The amino acid species of #1 to #12 are shown in capital single letter, with the metal uptake characteristics: +, transport; -, no transport; ?, promiscuous; /, not confirmed (a: Curie et al., 2000, b: Takahashi et al., 2011, c: Tiwari et al., 2014, d: Chang et al., 2020, e: Ishimaru et al., 2012, f: Ishikawa et al., 2012, g: Sasaki et al., 2012, h: Sui et al., 2018, i: Peris-Peris et al., 2017, j: Xia et al., 2010, k: Li et al., 2022, l: Yamaji et al., 2013, m: Yang et al., 2013, n: Thomine et al., 2000, o: Cailliatte et al., 2009, p: Li et al., 2019, q: Pottier et al., 2015, r: Alejandro et al., 2017, s: Gao et al., 2018, t: Zhao et al., 2018, u: Guo et al., 2022, v: Li et al., 2023, w: Ehrnstorfer et al., 2014, x: Bozzi et al., 2016a, y: Bozzi et al., 2016b).

In this study, we engineered the Mn/Cd uptake efficiencies of OsNramp5 by incorporating amino acid substitution(s) at the putative metal-binding residue(s) to produce rice plants that reduce Cd uptake efficiencies but retain that of Mn. The Mn/Cd uptake efficiency of the mutant OsNramp5 was first evaluated using a yeast model assay system (Podar et al., 2012; Chen et al., 2016). Together with further evaluation by direct measurement of the Mn/Cd uptake efficiencies, we identified several OsNramp5 mutants that exhibited desirable Mn/Cd uptake characteristics and retained more than 64.7% Mn uptake but reduced Cd uptake efficiency to background levels under the conditions mimicking a polluted soil. The molecular mechanisms of Mn/Cd uptake by OsNramp5 and mutant proteins are also discussed based on computational structural modeling.

## 2 Materials and Methods

### 2.1 Sequence alignment and phylogenetic analysis

The amino acid sequences of the 21 Nramps used in this study were obtained from the NCBI database (https://www.ncbi.nlm.nih.gov) and are listed in Supplementary Table 1, along with their NCBI accession numbers. Protein sequence alignment and phylogenetic analyses were conducted using MUSCLE (Edgar, 2004) and the Maximum likelihood method (Jones et al., 1992), respectively, in the MEGA 11 package (Tamura et al., 2021).

### 2.2 Plasmid construction of OsNramp5 and the mutants

The DNA sequence of the *OsNramp5* coding region was chemically synthesized by Geneart (Thermo Fisher Scientific, Waltham, MA, USA). The synthesized DNA was used as a template for polymerase chain reaction amplification using PrimeSTAR Max (Takara Bio, Shiga, Japan) and primers OsNramp5_GA_F and OsNramp5_GA_R. The polymerase chain reaction product was integrated into the expression plasmid of pDR195 (Rentsch et al., 1995; Addgene #36028), using the NEBuilder HiFi DNA Assembly Cloning Kit (New England Biolabs, Ipswich, MA, USA). The resulting plasmid was used for site-directed mutagenesis of *OsNramp5* using the primers listed in Supplementary Table 2. The DNA sequence was confirmed using Sanger sequencing.

### 2.3 Analysis of metal uptake characteristics of OsNramp5 mutants using yeast assay system

The yeast strains used in this study, BY4741(wild type; WT), Cd-sensitive *Δycf1* (Y04069), and Mn-sensitive *Δpmr1* (Y04534), were obtained from EUROSCARF (Oberursel, Germany). Plasmids containing the *OsNramp5* gene or mutants were introduced into yeast using YNB-ura (MP Biomedicals, Irvine, CA, USA) agar plates and the LiAc/SS carrier DNA/PEG method (Gietz and Schiestl, 2007). The obtained transformants were pre-cultured in liquid YNB-ura medium at 30 ℃, overnight with gentle shaking. The pre-cultured yeast was replaced with a new YNB-ura liquid medium by adjusting pH 4.5 and an optical density at 600 nm (OD600) = 0.001 in a final volume of 200 µl. CdSO_4_ was added to be 0 µM to 81.8 µM for the Cd sensitivity test.

The Cd sensitivity test was conducted using Cd-sensitive *Δycf1* by incubating the yeast in 96 plates with rubber lids at 30 °C for 3 days with gentle shaking. After incubation, OD600 was measured using EnSight Multimode Plate Reader (Perkin Elmer, Shelton, CT, USA) and IC_50_ values were calculated using R package “drc ver. 3.0-1” (Ritz et al., 2015). The relative Cd uptake efficiency was estimated using three biological replicates, regarding the reciprocal IC_50_ (1/IC_50_) of the empty plasmid and WT as 0% and 100%, respectively. The Mn sensitivity test was conducted using Mn-sensitive *Δpmr1* and a medium containing 0 mM to 1.5 mM MnSO_4_.

### 2.4 Quantification of the metal uptake characteristics using inductively coupled plasma optical emission spectroscopy (ICP-OES)

The WT yeast strain (BY4741) containing empty, *OsNramp5*, or the mutant plasmid was incubated in liquid YNB-ura medium at 30 ℃ overnight with gentle shaking. The pre-culture medium was transferred to 5 ml YNB-ura liquid medium containing 20 µM MnSO_4_, CdSO_4_, FeSO_4_, CuSO_4_, ZnSO_4_, and CoSO_4_, and adjusted to pH 4.5 and OD600 = 0.1. After 16 hours of incubation at 30 ℃ with gentle shaking, the cells were harvested via centrifugation, washed once with 20 mM EDTA, and washed three times with deionized water. After drying up the cell at 60 ℃ for three days, the cell was lysed by 60% HNO_3_. The concentrations of Cd and Mn in the cell lysates were measured using an Agilent 5800 inductively coupled plasma optical emission spectroscopy (ICP-OES; Agilent, Santa Clara, CA, USA) according to the manufacturer’s instructions. The apparent metal uptake was estimated from the actual metal uptake by subtracting the value for the empty plasmid from that of the background. The relative uptake value was calculated using the apparent metal uptake of the empty plasmid and WT as 0% and 100%, respectively. The Cd and Mn uptake efficiencies were analyzed at various Cd concentrations (Fig. 4) using a medium containing 1.8 mM MnSO_4_ and 0, 2.5, or 7.5 µM CdSO_4_. The relative uptake values were calculated using the apparent metal uptake of the empty plasmid and WT as 0% and 100%, respectively, for each CdSO_4_ concentration.

### 2.5 Computational structural analysis

The 3D structures of OsNramp5 and the mutants were predicted by AlphaFold2 running on the ColabFold v1.5.5 using MMseqs2 server (Mirdita et al., 2019, 2022; Jumper et al., 2021). The highest-ranking model was relaxed by using amber force fields (Eastman et al., 2017). The X-ray crystal structure of *Deinococcus radiodurans* Nramp (DraNramp)-G223W (Protein Data Bank ID: 8E6N) was chosen as the template for structural modeling (Ray et al., 2023). AlphaFill was used to predict the optimized conformation by incorporating Mn ions into the metal-binding sites of the predicted 3D structures (Hekkelman et al., 2023). Tunnel detection was performed by setting the tunnel bottleneck (minimum radius) to 0.9 Å, with the tunnel search starting point at approximately 3 Å from the Mn ion in CAVER Analyst 2.0 BETA (Jurcik et al., 2018), and the length and radius of the tunnels were calculated. Detection of cavities around the metal-binding sites was performed using pyKVFinder (Eisenberg et al., 1984; Guerra et al., 2021, 2023), as well as hydropathy calculations. The cavity detection was performed with the following settings: 0.9 Å probe size, within 3.5 Å to the direction of the metal transport tunnel entrance from Mn, and 3 Å from the center line of the transport tunnel. The predicted 3D structures were visualized using PyMOL v2.5.0 (https://pymol.org/).

## 3 Results

### 3.1 Sequence analysis of Nramps

To identify the putative amino acids of plant Nramps involved in metal binding, the amino acid sequence of the OsNramp5 protein was aligned and analyzed phylogenetically, with other 18 Nramps in monocot and dicot plants, as well as bacterial Nramps of *Staphylococcus capitis* Nramp (ScaNramp) and DraNramp (Fig. 1B and Supplementary Fig. 1). The key amino acid residues for the metal transporters were designated #1 to #12. Residues #2, #5, #8, and #11 were identified as metal-contacting residues based on structural analyses of bacterial Nramps (Ehrnstorfer et al., 2014). By focusing on these residues, aspartic acid (D) and asparagine (N) at #2 and #5, respectively, in TM1 were found to be highly conserved among all Nramps from bacteria to plants. In contrast, the metal-binding amino acids at positions #8 and #11 in TM6 are diverse. All Nramps transport Cd, which has been characterized so far, containing methionine (M) at position #11 (OsNramp1, 2, and 5; *Arabidopsis thaliana* Nramp (AtNramp) 1, 3, 4, and 6; *Zea mays* Nramp (ZmNramp) 2 and 5; ScaNramp; and DraNramp). Nramp, which has been characterized as not transporting Cd, contains different amino acids at #11, such as valine (V) or threonine (T) (OsNramp6, OsNramp4 (also known as OsNrat1), and ZmNrat1). Threonine (T) at position #11 is associated with unique metal uptake characteristics of the trivalent metal ion Al in OsNramp4 and ZmNrat1. No clear relationship was found between the amino acid species at position #8 and the metal uptake characteristics.

These observations suggest that the amino acids in TM1 (#2 and #5) are indispensable for their crucial function as transporters and that amino acids #8 and #11 in TM6 may be involved in metal uptake selectivity. This hypothesis is supported by previous studies on the metal selectivity modification of amino acid substitutions at position #11 in several Nramps (Bozzi et al., 2016a; Li et al., 2018; Lu et al., 2018). Therefore, we modified residues #8 and #11 to engineer OsNramp5 to reduce Cd uptake efficiency.

### 3.2 Analysis of metal uptake characteristics of OsNramp5 mutants using yeast assay system

The yeast assay system is a well-established system for estimating the metal uptake characteristics of transporter proteins of interest. Metal-sensitive mutant strains of yeast show retarded growth in the presence of a particular metal ion (Podar et al., 2012; Chen et al., 2016; Li et al., 2018). To evaluate the function of residue #11 of OsNramp5, corresponding to 235^th^ Met, the residue was substituted with other 19 amino acid species. The obtained OsNramp5 mutants, designated as M235X, were transformed to the Cd-sensitive yeast strain (*Δycf1*; Y04069), together with the previously reported mutants of Q337K and A512T (Kuramata et al., 2022; Qu and Nakanishi, 2023). The growth of Cd-sensitive yeast at various Cd concentrations was measured using OD600 to calculate IC_50_. The IC_50_ value was used to estimate the metal uptake efficiency (Supplementary Fig. 2). All 19 mutant proteins constructed in this study exhibited significantly lower Cd uptake (less than 20%) than that of WT (Fig. 2A). Mn uptake efficiency was also analyzed because OsNramp5 is responsible for Mn uptake, which is an essential micronutrient that facilitates the growth and development of plants. The Mn uptake efficiencies were highly variable among the mutants when the assay was conducted using the Mn-sensitive yeast strain (*Δpmr1*; Y04534). However, the substitution to alanine (A) and cysteine (C) exhibited more than 60% Mn transport compared to WT (M235A and M235C; Fig. 2B). The Mn/Cd uptake ratio was calculated by dividing the relative Mn uptake by the relative Cd uptake, to evaluate the metal selectivity of the OsNramp5 mutants (Table 1). The Mn/Cd uptake ratios of M235A and M235C were 11.3 and 7.5, respectively, indicating that these mutants had the expected characteristics of low Cd uptake while retaining Mn uptake. The Mn/Cd uptake ratio of the previously characterized Q337K allele was estimated as 1.0. The Q337K allele transported both Cd and Mn at a medium level compared with WT (36.0% and 37.2%, respectively). This is consistent with a previous report showing that the Q337K allele does not affect the selectivity of Cd or Mn transport (Kuramata et al., 2022). Another previously characterized mutant, A512T, also showed a Mn/Cd uptake ratio of 0.9, due to an approximately 50% reduction in both Mn and Cd uptake. This result was in contrast to that of a previous study in which the mutant showed only a reduction in Cd uptake in yeast (Qu and Nakanishi, 2023). Taken together, M235A and M235C showed the expected metal uptake characteristics, that is, little Cd uptake and efficient Mn uptake, in the yeast assay system; therefore, these mutants were used for subsequent studies.

**Figure 2.**
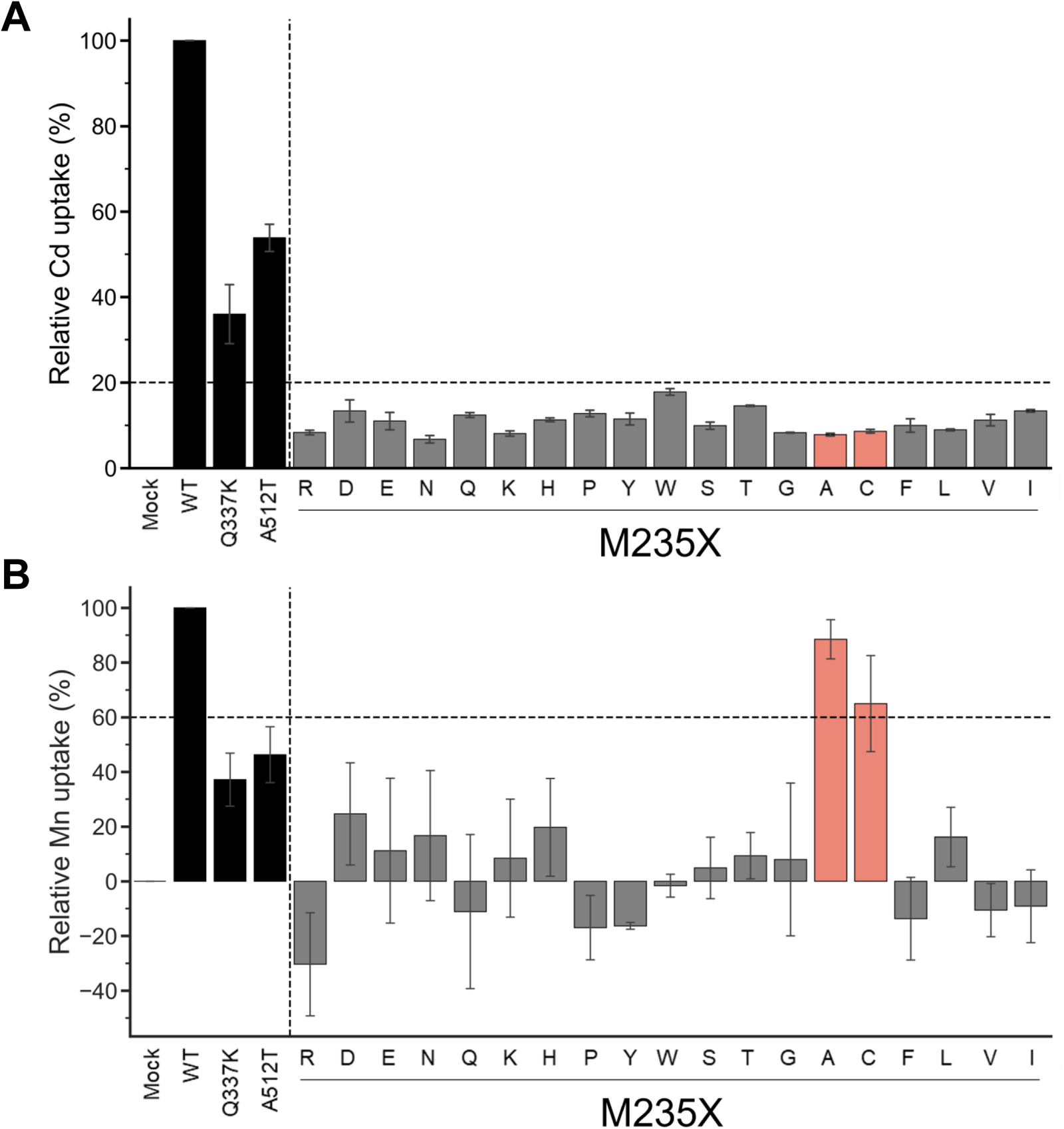
Cd and Mn uptake efficiencies of M235X mutants of OsNramp5. **(A)** Cd uptake efficiency analyzed for 19 M235X mutants in the yeast assay system, together with the previously reported mutants of Q337K and A512T. The mean and standard error of relative Cd uptake efficiency are shown (N=3, WT=100%). The mutants exhibited <20% Cd and >60% Mn uptake efficiencies are highlighted in red. **(C)** Mn uptake efficiencies are shown as in (B).

**Table 1.** Relative Cd and Mn uptake efficiency and Mn/Cd ratio of OsNramp5 M235X mutants.

| Construct | Mn/Cd ratio | Relative Cd uptake (%) |  |  |  | Relative Mn uptake (%) |  |  |  |
| --- | --- | --- | --- | --- | --- | --- | --- | --- | --- |
|  |  | Mean | ± | SE | *p*-value | Mean | ± | SE | *p*-value |
| Mock | - | 0.0 | ± | 0.0 | ** | 0.0 | ± | 0.0 | ** |
| WT | 1.0 | 100.0 | ± | 0.0 |  | 100.0 | ± | 0.0 |  |
| Q337K | 1.0 | 36.0 | ± | 6.9 | ** | 37.2 | ± | 9.7 | ** |
| A512T | 0.9 | 53.9 | ± | 3.2 | ** | 46.3 | ± | 10.2 | ** |
| M235A | 11.3 | 7.8 | ± | 0.3 | ** | 88.5 | ± | 7.2 |  |
| M235C | 7.5 | 8.6 | ± | 0.4 | ** | 65.0 | ± | 17.6 |  |
| M235N | 2.5 | 6.7 | ± | 0.9 | ** | 16.7 | ± | 23.8 | ** |
| M235D | 1.8 | 13.4 | ± | 2.6 | ** | 24.6 | ± | 18.7 | ** |
| M235L | 1.8 | 8.9 | ± | 0.2 | ** | 16.2 | ± | 10.9 | ** |
| M235H | 1.7 | 11.3 | ± | 0.5 | ** | 19.7 | ± | 17.8 | ** |
| M235K | 1.0 | 8.1 | ± | 0.6 | ** | 8.5 | ± | 21.6 | ** |
| M235E | 1.0 | 11.0 | ± | 2.0 | ** | 11.2 | ± | 26.5 | ** |
| M235G | 1.0 | 8.3 | ± | 0.1 | ** | 8.0 | ± | 28.0 | ** |
| M235T | 0.6 | 14.6 | ± | 0.2 | ** | 9.3 | ± | 8.5 | ** |
| M235S | 0.5 | 9.9 | ± | 0.9 | ** | 4.9 | ± | 11.2 | ** |
| M235W | -0.1 | 17.8 | ± | 0.8 | ** | -1.6 | ± | 4.2 | ** |
| M235I | -0.7 | 13.4 | ± | 0.3 | ** | -9.1 | ± | 13.3 | ** |
| M235Q | -0.9 | 12.4 | ± | 0.6 | ** | -11.1 | ± | 28.2 | ** |
| M235V | -0.9 | 11.2 | ± | 1.3 | ** | -10.5 | ± | 9.8 | ** |
| M235P | -1.3 | 12.8 | ± | 0.8 | ** | -17.0 | ± | 11.8 | ** |
| M235F | -1.4 | 10.0 | ± | 1.6 | ** | -13.7 | ± | 15.1 | ** |
| M235Y | -1.4 | 11.5 | ± | 1.4 | ** | -16.3 | ± | 1.2 | ** |
| M235R | -3.7 | 8.3 | ± | 0.5 | ** | -30.4 | ± | 18.9 | ** |

### 3.3 Engineering of OsNramp5 by a double mutation

Amino acid substitutions focusing on residue #11 yielded the OsNramp5 mutants, M235A and M235C, with improved Mn/Cd uptake characteristics. The mutants were further subjected to amino acid substitution at residue #8, another putative residue involved in metal uptake selectivity (Fig. 1B). Residue #8 for OsNramp5, corresponding to the 232^nd^ Ala, was substituted with cysteine, serine, or valine; these amino acid species are found at residue #8 in other Nramp families. The 232^nd^ Ala residue was also substituted with methionine, which may play a significant role in Cd and Mn uptake at residue #11 (235^th^ Met in OsNramp5). The metal uptake characteristics of the obtained double mutants were analyzed in a yeast assay system, together with mutants containing a single amino acid substitution at position 232. All double mutants showed less than 20% Cd uptake efficiency compared to WT (Fig. 3A). When the Mn uptake efficiencies were examined, three double mutants, A232C+M235A, A232S+M235A, and A232S+M235C, showed Mn uptake efficiencies more than 60% (Fig. 3B). Two single mutants (A232M and A232V) showed substantially lower Cd uptake efficiencies, although Mn uptake efficiencies were also impaired. Interestingly, the A232M+M235A mutant was simply a swap of the amino acid species between positions 232 and 235 from the WT. However, A232M+M235A displayed extremely low uptake efficiencies for both Cd and Mn, suggesting that both the position and amino acid species determine the efficiency of Cd/Mn uptake.

**Figure 3.**
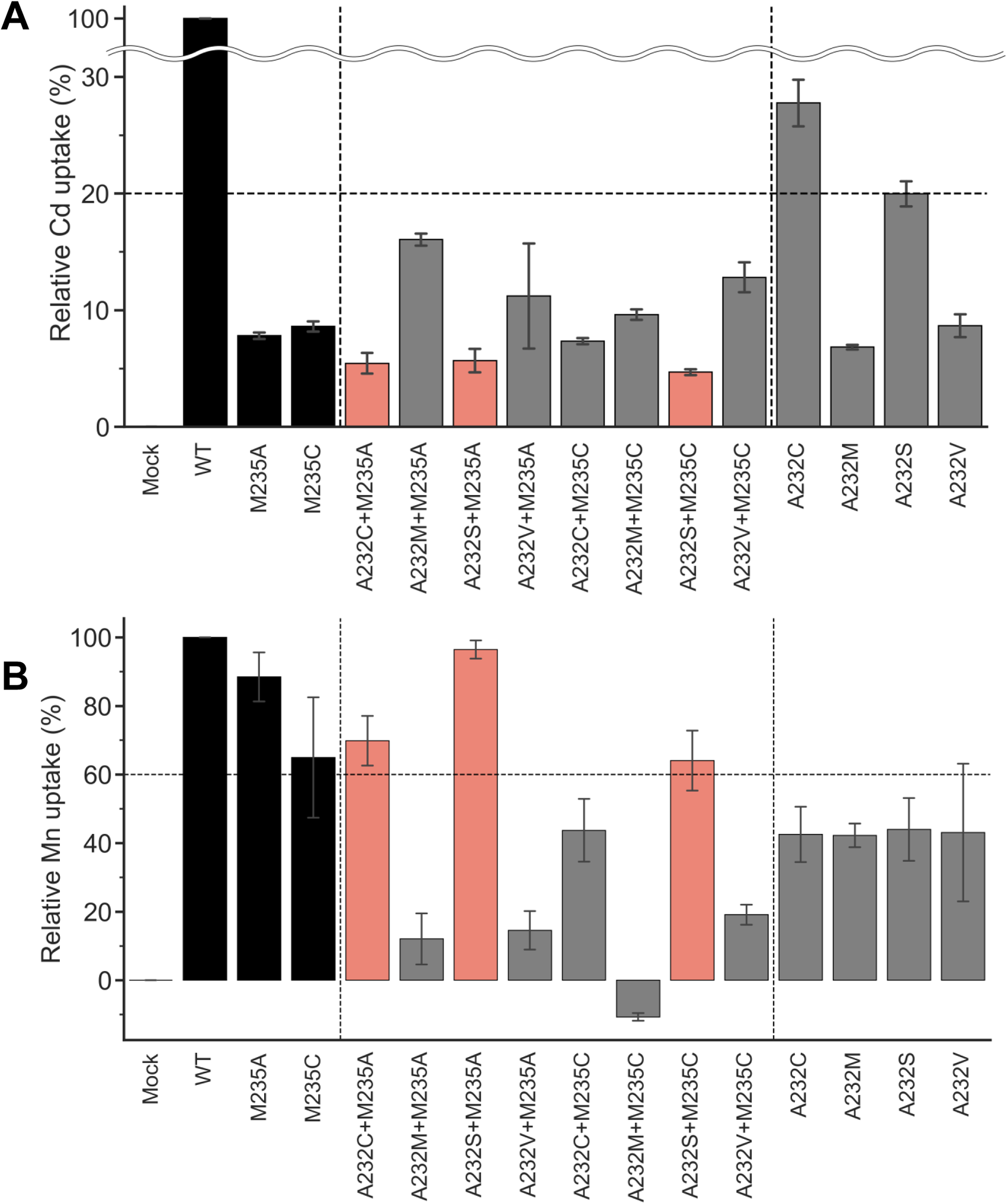
Cd and Mn uptake efficiencies of OsNramp5 double mutants. **(A)** Cd uptake efficiency analyzed for eight double mutants in the yeast assay system, together with four single mutants at residue 232. M235A and M235C were also used as references. The mean and standard error of relative Cd uptake efficiency are shown (N=3, WT=100%). The mutants exhibited <20% Cd and >60% Mn uptake efficiencies are highlighted in red. **(B)** Mn uptake efficiencies are shown as in (A).

The Mn/Cd uptake ratios of the double mutants were estimated for the single M235X mutants (Table 2). Three double mutants, A232S+M235A, A232S+M235C, and A232C+M235A, showed better Mn/Cd uptake ratios of 17.0, 13.6, and 12.8, respectively, than the single mutants (11.3, M235A; 7.5, M235C). Notably, A232S+M235A showed the most desirable metal uptake characteristics among the mutants in this study, as shown by the Mn/Cd uptake ratio of 17.0, with an equivalent Mn uptake efficiency (96.4%) compared to that of WT, although the Cd uptake efficiency was significantly reduced (5.7%) (Fig. 3A, Table 2).

**Table 2.** Relative Cd and Mn uptake efficiency and Mn/Cd ratio of OsNramp5 single and double mutants.

| Construct | Mn/Cd ratio | Relative Cd uptake (%) |  |  |  | Relative Mn uptake (%) |  |  |  |
| --- | --- | --- | --- | --- | --- | --- | --- | --- | --- |
|  |  | Mean | ± | SE | *p*-value | Mean | ± | SE | *p*-value |
| Mock | - | 0.0 | ± | 0.0 | ** | 0.0 | ± | 0.0 | ** |
| WT | 1.0 | 100.0 | ± | 0.0 | ** | 100.0 | ± | 0.0 |  |
| M235A | 11.3 | 7.8 | ± | 0.3 |  | 88.5 | ± | 7.2 |  |
| M235C | 7.5 | 8.6 | ± | 0.4 |  | 65.0 | ± | 17.6 |  |
| A232S+M235A | 17.0 | 5.7 | ± | 1.0 |  | 96.4 | ± | 2.6 |  |
| A232S+M235C | 13.6 | 4.7 | ± | 0.2 |  | 64.1 | ± | 7.1 |  |
| A232C+M235A | 12.8 | 5.5 | ± | 0.9 |  | 69.8 | ± | 7.2 |  |
| A232M | 6.2 | 6.8 | ± | 0.2 |  | 42.3 | ± | 3.5 | ** |
| A232C+M235C | 6.0 | 7.3 | ± | 0.3 |  | 43.7 | ± | 9.1 | ** |
| A232V | 5.0 | 8.7 | ± | 1.0 |  | 43.1 | ± | 20.1 | ** |
| A232S | 2.2 | 20.0 | ± | 1.1 | ** | 44.0 | ± | 9.1 | ** |
| A232C | 1.5 | 27.8 | ± | 2.0 | ** | 42.5 | ± | 8.0 | ** |
| A232V+M235C | 1.5 | 12.8 | ± | 1.3 | ** | 19.1 | ± | 2.9 | ** |
| A232V+M235A | 1.3 | 11.2 | ± | 4.5 |  | 14.5 | ± | 6.5 | ** |
| A232M+M235A | 0.8 | 16.1 | ± | 0.5 | ** | 12.1 | ± | 7.4 | ** |
| A232M+M235C | -1.1 | 9.6 | ± | 0.5 |  | -10.7 | ± | 1.1 | ** |

### 3.4 Direct measurement of the metal uptake of OsNramp5 mutants in yeast

The yeast assay system is a powerful tool that enables high-throughput screening of various mutants. However, the evaluation relies on the growth of yeast, which may be affected by various inter-/intra-cellular conditions. Therefore, the actual metal uptake efficiencies of the five selected mutants (M235A, M235C, A232C+M235A, A232S+M235A, and A232S+M235C) were directly determined using inductively coupled plasma optical emission spectrometry (ICP-OES). The mutants were transformed into the WT yeast strain and cultured overnight in the presence of 20 µM Cd and 20 µM Mn. The incorporated metal content was determined by ICP-OES.

ICP-OES analysis showed that the Cd uptake of the five mutants was lower than that of the WT, coinciding with the results of the yeast assay system (Table 3). The relative Cd uptake efficiencies were estimated to range from 10.9% to 22.3% against that of WT. When the Mn uptake was examined, the mutants displayed similar or decreased Mn uptake efficiencies compared to the WT, as shown by the relative values from 54.8% to 103.3% (Table 3). The best Mn/Cd ratio of 7.4 was raised by A232C+M235A. Cd uptake was reduced to 13.9% of WT, with no influence on Mn uptake efficiency (103.3%) (Table 3). The Mn/Cd ratio of the single mutant M235A was 3.5, suggesting that the double mutation at residues #8 and #11 successfully improved metal uptake characteristics.

**Table 3.** Direct measurement of Cd and Mn contents for selected OsNramp5 mutants.

| Construct | Mn/Cd ratio | Cd uptake |  |  |  |  | Mn uptake |  |  |  |  |
| --- | --- | --- | --- | --- | --- | --- | --- | --- | --- | --- | --- |
|  |  | Relative value (%) | Mean (ng/mg) | ± | SE | p-value | Relative value (%) | Mean (ng/mg) | ± | SE | p-value |
| Mock | - | 0.0 | 15.5 | ± | 1.1 | ** | 0.0 | 7.5 | ± | 0.4 | ** |
| WT | 1.0 | 100.0 | 129.9 | ± | 5.8 |  | 100.0 | 17.0 | ± | 1.4 |  |
| A512T | 0.5 | 90.4 | 118.9 | ± | 11.3 |  | 48.7 | 12.2 | ± | 0.9 | ** |
| M235A | 3.5 | 17.2 | 35.3 | ± | 2.0 | ** | 60.2 | 13.2 | ± | 0.7 | * |
| M235C | 5.0 | 10.9 | 28.0 | ± | 1.9 | ** | 54.8 | 12.7 | ± | 0.6 | * |
| A232C+M235A | 7.4 | 13.9 | 31.4 | ± | 2.8 | ** | 103.3 | 17.3 | ± | 0.4 |  |
| A232S+M235A | 3.7 | 22.3 | 41.0 | ± | 2.9 | ** | 82.8 | 15.4 | ± | 0.5 |  |
| A232S+M235C | 4.6 | 13.5 | 31.0 | ± | 4.1 | ** | 62.5 | 13.5 | ± | 1.2 | * |

The Cd/Mn content of the yeast was measured using ICP-OES. The results were analyzed based on the relative Cd/Mn uptake (%) and Mn/Cd ratio. The mean and standard error (SE) are shown (N=3) with the statistical evaluation compared to the WT using Dunnett's test (\* $p < 0.05$ , \*\* $p < 0.01$ ).

A series of experiments using a yeast assay system and direct measurements by ICP-OES identified several desirable OsNramp5 mutants. Three mutants were selected for further characterization: M235A, the best single mutant in the yeast assay system (Table 1); A232S+M235A, the best double mutant in the yeast assay system (Table 2); and A232C+M235A, the best mutant in the ICP-OES analysis (Table 3).

### 3.5 Cd uptake characterization of the mutants under various Cd concentrations

In natural conditions, the average soil Cd concentration has been reported as 0.36 mg/kg globally, 0.27 mg/kg in the USA, 0.15 mg/kg in Europe, and 0.27 mg/kg in China (Hutchinson et al., 1987; Zhang et al., 2015; Kubier et al., 2019; Ballabio et al., 2023). In Japan, the soil Cd concentration was reported as an average of 0.30 mg/kg and a maximum of 0.80 mg/kg (Asami et al., 1988). Therefore, the Cd uptake of the representative OsNramp5 mutants (M235A, A232S+M235A, and, A232C+M235A) was examined under three conditions: 0 µM for the blank; 2.5 µM (≈ 0.28 mg/kg), mimicking the normal condition; and 7.5 µM (≈ 0.84 mg/kg), mimicking the highly polluted condition. The Mn concentration was determined at 1.8 mM (≈ 98.9 mg/kg) because the previous study of OsNramp5 knockout mutant showed that brown spot disease started to be observed at a soil concentration of 51.9 mg/kg Mn, but not of 220.8 mg/kg Mn (Honma et al., 2017).

This analysis showed that the Cd uptake of all the selected OsNramp5 mutants (A232S+M235A, A232C+M235A, and M235A) was estimated at background levels, even at high Cd concentrations (Fig. 5A). A232S+M235A, A232C+M235A, and M235A showed constant Mn uptakes under all Cd concentrations: 91.9% to 107.9% of WT in the absence of Cd, 79.8% to 84.0% of WT in the 2.5 µM Cd, and 64.7% to 76.1% of WT at the 7.5 µM Cd (Fig. 5B and Table 4). Altogether, this dose-dependent evaluation further strengthened the fact that the three mutants (M235A, A232C+M235A, and A232S+M235A) showed desirable Cd and Mn uptake characteristics under conditions mimicking polluted soil.

**Figure 4.**
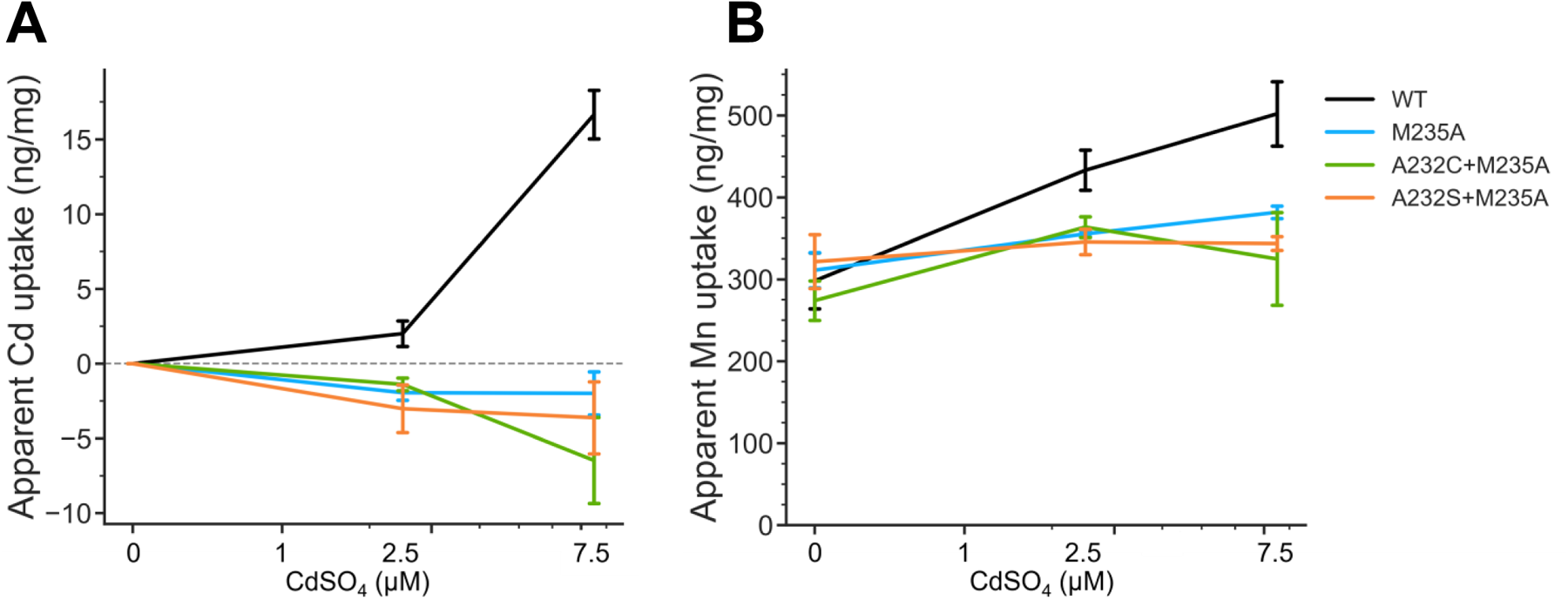
Cd and Mn uptake efficiency in various Cd concentrations. **(A)** Apparent Cd uptake amount measured using ICP-OES for the WT yeast strain containing OsNramp5 (WT) and the mutants (M235A, A232C+M235A, and A232S+M235A) in the presence of different Cd concentrations (0, 1, 2.5, and 7.5 μM). The mean and standard error were shown (N = 3). **(B)** Apparent Mn uptake amount measured in the same condition as (A).

**Figure 5.**
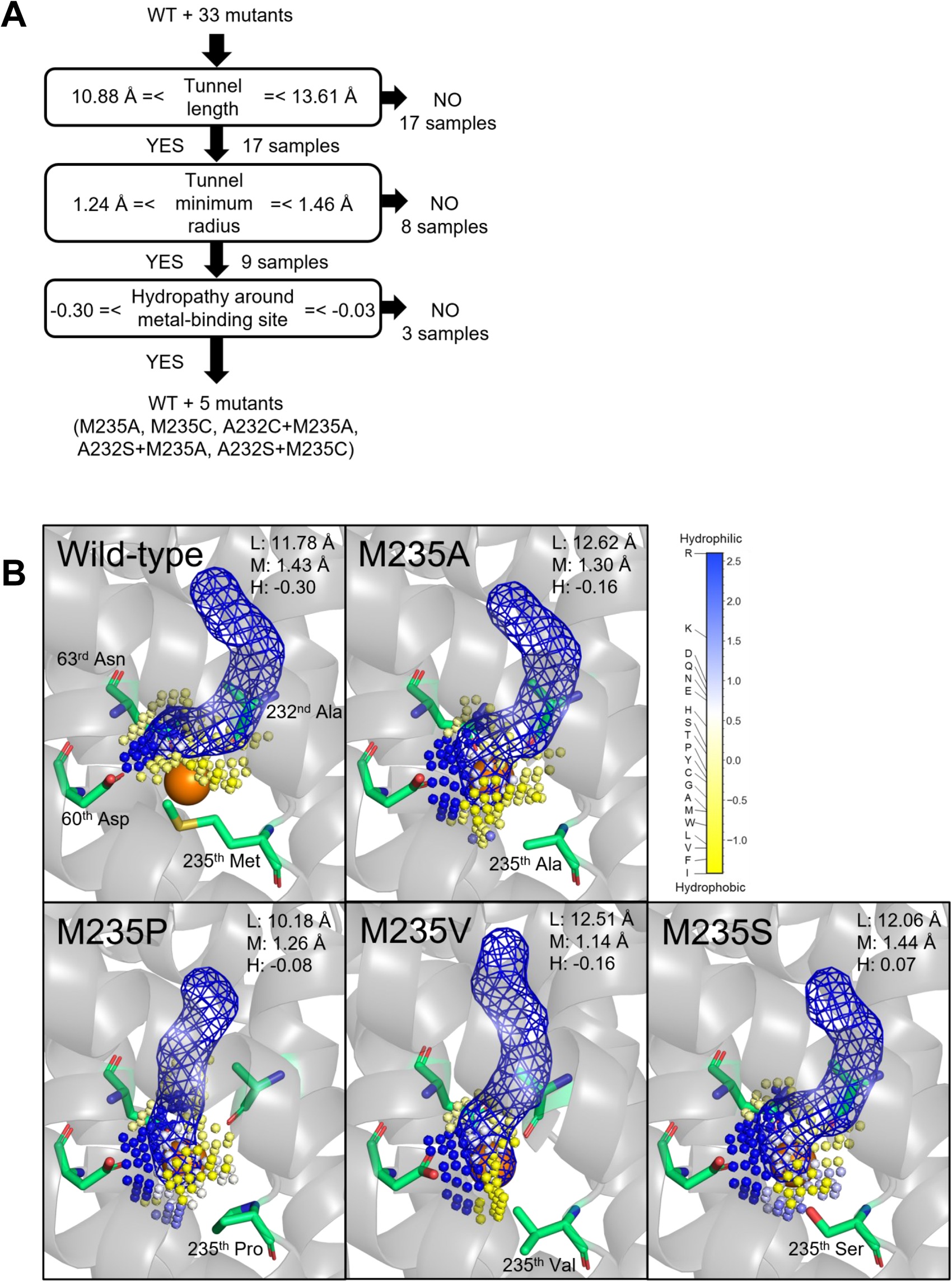
Structure and chemical properties that may determine the Cd and Mn uptake efficiency of OsNramp5. **(A)** Appropriate tunnel length, tunnel minimum radius, and hydropathy around the metal-binding site hypothesized to be required for efficient Mn uptake (>60% of WT). The number of mutants that passed in the node is shown. **(B)** Structure of the predicted transport tunnel and metal-binding site of WT, M235A, M235P, M235V, and M235S. The predicted tunnels are visualized by blue mesh, together with the key residues. The predicted cavity around the metal-binding site is shown by a small sphere, which is colored according to the Eisenberg & Weiss hydropathy scale. The properties of the length (L) and minimum radius (M) of the transport tunnel, and hydropathy around the metal-binding site (H), are shown.

**Table 4.** Apparent Cd and Mn uptake in various Cd concentrations for selected OsNramp5 mutants.

| Construct | CdSO<sub>4</sub> | Apparent Cd uptake (ng/mg) |  |  |  |  | Apparent Mn uptake (ng/mg) |  |  |  |  |
| --- | --- | --- | --- | --- | --- | --- | --- | --- | --- | --- | --- |
|  |  | Relative value (%) | Mean | ± | SE | p-value | Relative value (%) | Mean | ± | SE | p-value |
| Mock | 0 μM | - | N.D. | | | | 0.0 | 0.0 | ± | 0.0 | ** |
| | 2.5 μM | 0.0 | 0.0 | ± | 0.0 | | 0.0 | 0.0 | ± | 0.0 | ** |
| | 7.5 μM | 0.0 | 0.0 | ± | 0.0 | ** | 0.0 | 0.0 | ± | 0.0 | ** |
| WT | 0 μM | - | N.D. | | | | 100.0 | 298.0 | ± | 34.2 | |
| | 2.5 μM | 100.0 | 2.0 | ± | 0.8 | | 100.0 | 433.1 | ± | 24.6 | |
| | 7.5 μM | 100.0 | 16.7 | ± | 1.6 | | 100.0 | 501.9 | ± | 39.2 | |
| M235A | 0 μM | - | N.D. | | | | 104.4 | 311.1 | ± | 21.5 | |
| | 2.5 μM | 0 > | -1.9 | ± | 0.5 | * | 82.0 | 355.3 | ± | 3.4 | ** |
| | 7.5 μM | 0 > | -2.0 | ± | 1.4 | ** | 76.1 | 381.7 | ± | 7.7 | |
| A232C+M235A | 0 μM | - | N.D. | | | | 91.9 | 273.9 | ± | 24.0 | |
| | 2.5 μM | 0 > | -1.4 | ± | 0.4 | | 84.0 | 363.6 | ± | 12.6 | * |
| | 7.5 μM | 0 > | -6.5 | ± | 2.9 | ** | 64.7 | 324.8 | ± | 56.7 | ** |
| A232S+M235A | 0 μM | - | N.D. | | | | 107.9 | 321.5 | ± | 32.8 | |
| | 2.5 μM | 0 > | -3.0 | ± | 1.6 | ** | 79.8 | 345.6 | ± | 15.4 | ** |
| | 7.5 μM | 0 > | -3.6 | ± | 2.4 | ** | 68.5 | 343.7 | ± | 8.3 | * |

## 4 Discussion

Phylogenetic analysis and metal uptake characterization of OsNramp5 mutants using a yeast assay system and direct measurement of metal uptake by ICP-OES successfully identified several OsNramp5 mutants that effectively transported Mn but reduced Cd uptake efficiency compared to the original OsNramp5. The compatibility between the results of the yeast assay system and ICP-OES was evaluated by estimating the correlation coefficient. That was estimated as R=0.95 with *p* < 0.01 and R=0.85 with *p* < 0.01 for the Cd and Mn uptake, respectively (Supplementary Fig. 3), indicating that both experimental methods were sufficiently worked to evaluate metal uptake characteristics of OsNramp5 mutants. Notably, the Cd dose-dependent test showed that the three mutants (A232S+M235A, A232C+M235A, and M235A) retained more than 64.7% Mn uptake efficiency but background levels of Cd uptake in the conditions mimicking the polluted soil.

Phylogenetic analysis of OsNramp5 and previous studies suggested key residues of TM1 and TM6 for metal uptake efficiency and selectivity. Aspartic acid and asparagine are highly conserved at residues #2 and #5 of TM1, respectively, in various metal ion transporters and are presumably essential for their function as transporters (Fig.1; Chaloupka et al., 2005; Bozzi et al., 2020). Key residues #8 and #11 are degenerated in the Nramp family (Fig. 1). The involvement of residues #8 and #11 in metal uptake selectivity was partially demonstrated in this study. All amino acid substitutions at residue #11, M235X of OsNramp5, showed a significant reduction in Cd transport activity compared to the WT (<20%), suggesting that methionine at the metal-binding site is crucial for Cd transport. Mutations #8 and #11 in TM6 frequently reduced the Mn uptake efficiency, whereas efficient Mn transport was achieved when methionine (WT), alanine, and cysteine were positioned at either #8 or #11 (Fig. 2B & 3 B).

To gain mechanistic insights into the Cd/Mn uptake efficiencies of the OsNramp5 mutants analyzed in this study, their 3D structures were computationally predicted. Several amino acids surrounding the predicted metal transport tunnel of the original OsNramp5 have been reported to affect the metal selectivity of various Nramps (73^rd^ Asn, 225^th^ Asp, 228^th^ Ala, 393^th^ Ile, and 396^th^ Ser; Supplementary Fig. 4) (Pottier et al., 2015; Bozzi et al., 2019; Sun et al., 2019), suggesting the reliability of the predicted structure. The predicted structure suggested several requirements that may determine the Cd/Mn uptake efficiencies: the tunnel length, minimum radius as a tunnel width, and hydropathy around the metal-binding sites of the predicted structures.

For the high Mn uptake efficiency (>60%), the first requirement was supposed to be tunnel length, as shown that the mutants showed more than 60% Mn uptake efficiency have tunnel lengths between 10.88 and 13.61 Å (Fig. 5A and Supplementary Figure 5). In this first node, 17 proteins were eliminated from 34 samples (WT and 33 mutants). A typical example of a change in the tunnel length and shape was observed in the predicted structure of M235P, whose tunnel length was predicted as 10.18 Å (Fig. 5B). Proline substitutions are not easily accommodated in transmembrane helices, especially near the middle of the helix (Yohannan et al., 2004). The second requirement was suggested to be a tunnel minimum radius between 1.24 and 1.46 Å. Eight out of 17 proteins were eliminated by this step, as shown in the typical example of the predicted structure of M235V with the tunnel minimum radius of 1.14 Å (Fig. 5B). The third requirement is hypothesized to be the hydrophilicity of the metal-binding sites. The mutants that showed more than 60% Mn uptake efficiency have hydrophilicity between - 0.30 and -0.03, and six proteins (WT and five mutants) were passed in this step. It is presumed that an appropriate hydrophilicity is required to incorporate metal ions after they pass through the transport tunnel.

For the Cd uptake efficiency, methionine at residue #11 was critical for Cd transport. All amino acid substitutions of residue #11 and M235X of OsNramp5 showed a significant reduction of Cd transport efficiency compared to the WT (<20%). The crucial role of methionine at residue #11 has been demonstrated in other Nramp (Bozzi et al., 2016a). The necessity of methionine for Cd uptake has been discussed in a previous report, which provided the significant stabilization necessary for the binding and transport of Cd, which can forge strong covalent-like interactions with sulfur (Bozzi et al., 2016a). The low to moderate Cd uptake efficiency due to the mutation of other positions (A232C, Q337K, and A512T) can be explained by the requirement of tunnel length and minimum radius for efficient Mn uptake. We speculate that these three proteins did not allow Cd ions to reach the metal-binding site because their metal transport tunnels did not match the requirements of either or both spatial properties for metal transport (Supplementary Figure 5). In brief, structural prediction and mutant analysis in the yeast assay system suggested that metal uptake efficiency may be determined by the tunnel length, tunnel minimum radius, and hydropathy around the metal-binding site. It is interesting to see these observations can be applied to the semi-rational engineering of other Nramps.

This study identified several OsNramp5 mutants that showed desirable metal uptake efficiencies in yeast, including high Mn uptake and low Cd uptake, even under various Mn and Cd concentrations that mimic natural soil conditions (Table 3 and Fig. 4). However, it is indispensable to evaluate Cd/Mn uptake characteristics in plants. The metal uptake characteristics of OsNramp5-Q337K have been evaluated in rice, and the Cd and Mn contents in the straw of the mutant rice were estimated as 40% of those in WT rice (Kuramata et al., 2022). This is a comparable value with the estimated Cd and Mn uptake efficiency in this study (40%; Fig. 2). In another study, it was reported that in the analysis of amino acid substitution mutants of AtNramp1, Mn concentrations in yeast showed the same tendency as in plants results (Fu et al., 2022). These observations suggest that the characteristics of the OsNramp5 mutant can also be observed in rice plants. The obtained three mutants (M235A, A232C+M235A, and A232S+M235A) which showed Cd uptake of background level even in high concentrations of Cd (7.5 µM; Fig. 4) should be the priorities for the evaluation in rice plants.

Future studies should apply a prime editor technique to incorporate M235A, A232C+M235A, or A232S+M235A mutations *in planta* (Anzalone et al., 2019). This technique enables precise base substitution and eventually produces a final product with no foreign genes in the genome. Several prime editing applications have been reported in various plants (Butt et al., 2020; Lin et al., 2020; Zong et al., 2022). The double mutation in this study can be managed by a single prime editing step, as shown by mutations at two positions of DNA 40 nucleotides apart (Tao et al., 2022).

## Author Contributions

Experimental Design, T.N. and J.I.; Performed experiments, J.I.; Data evaluation, T.N., J.I., T.T., and T.K.; Writing, T.N. and J. I., with support from T.T. and T.K.

## Acknowledgments

We would like to thank all our colleagues at Nakamura’s laboratory at Kyushu University for their kind advice regarding the experimental protocol and discussions. The ICP-OES (Agilent 5800) measurements were performed at the Center for Advanced Instrumental and Educational Support, Faculty of Agriculture, Kyushu University, Japan.

## References

Alejandro, S., Cailliatte, R., Alcon, C., Dirick, L., Domergue, F., Correia, D., et al. (2017). Intracellular distribution of manganese by the trans-golgi network transporter NRAMP2 is critical for photosynthesis and cellular Redox homeostasis. Plant Cell 29, 3068–3084. doi: 10.1105/tpc.17.00578

Anzalone, A. V., Randolph, P. B., Davis, J. R., Sousa, A. A., Koblan, L. W., Levy, J. M., et al. (2019). Search-and-replace genome editing without double-strand breaks or donor DNA. Nature 576, 149–157. doi: 10.1038/s41586-019-1711-4

Asami, T., Kubota, M., and Minamisawa, K. (1988). Natural Abundance of Cadmium, Antimony, Bismuth and Some Other Heavy Metals in Japanese Soils. Soil Sci. Plant. Nutr. 59, 197–199. doi: 10.20710/dojo.59.2_197

Ballabio, C., Jones, A., Montanarella, L., and Toth, G. (2023). Cadmium in the Soils of the EU Analysis of LUCAS Soils data for the review of Fertilizer Directive., in JRC Technical Reports. doi: 10.2760/3942

Bozzi, A. T., Bane, L. B., Weihofen, W. A., McCabe, A. L., Singharoy, A., Chipot, C. J., et al. (2016a). Conserved methionine dictates substrate preference in Nramp-family divalent metal transporters. Proc. Natl. Acad. Sci. U S A 113, 10310–10315. doi: 10.1073/pnas.1607734113

Bozzi, A. T., Bane, L. B., Weihofen, W. A., Singharoy, A., Guillen, E. R., Ploegh, H. L., et al. (2016b). Crystal Structure and Conformational Change Mechanism of a Bacterial Nramp-Family Divalent Metal Transporter. Structure 24, 2102–2114. doi: 10.1016/j.str.2016.09.017

Bozzi, A. T., Zimanyi, C. M., Nicoludis, J. M., Lee, B. K., Zhang, C. H., and Gaudet, R. (2019). Structures in multiple conformations reveal distinct transition metal and proton pathways in an Nramp transporter. Elife 8. doi: 10.7554/eLife.41124.001

Bozzi, A. T., McCabe, A. L., Barnett, B. C., and Gaudet, R. (2020). Transmembrane helix 6b links proton and metal release pathways and drives conformational change in an Nrampfamily transition metal transporter. J. Biol. Chem. 295, 1212–1224. doi: 10.1074/jbc.RA119.011336

Bozzi, A. T., and Gaudet, R. (2021). Molecular Mechanism of Nramp-Family Transition Metal Transport. J. Mol. Biol. 433. doi: 10.1016/j.jmb.2021.166991

Butt, H., Rao, G. S., Sedeek, K., Aman, R., Kamel, R., and Mahfouz, M. (2020). Engineering herbicide resistance via prime editing in rice. Plant Biotechnol. J 18, 2370–2372. doi: 10.1111/pbi.13399

Cailliatte, R., Lapeyre, B., Briat, J. F., Mari, S., and Curie, C. (2009). The NRAMP6 metal transporter contributes to cadmium toxicity. Biochemical Journal 422, 217–228. doi: 10.1042/BJ20090655

Cailliatte, R., Schikora, A., Briat, J. F., Mari, S., and Curie, C. (2010). High-affinity manganese uptake by the metal transporter nramp1 is essential for Arabidopsis growth in low manganese conditions. Plant Cell 22, 904–917. doi: 10.1105/tpc.109.073023

Chaloupka, R., Courville, P., Veyrier, F., Knudsen, B., Tompkins, T. A., and Cellier, M. F. M. (2005). Identification of functional amino acids in the Nramp family by a combination of evolutionary analysis and biophysical studies of metal and proton cotransport in vivo. Biochemistry 44, 726–733. doi: 10.1021/bi048014v

Chang, J. D., Huang, S., Yamaji, N., Zhang, W., Ma, J. F., and Zhao, F. J. (2020). OsNRAMP1 transporter contributes to cadmium and manganese uptake in rice. Plant Cell. Environ. 43, 2476–2491. doi: 10.1111/pce.13843

Chen, X., Li, J., Wang, L., Ma, G., and Zhang, W. (2016). A mutagenic study identifying critical residues for the structure and function of rice manganese transporter OsMTP8.1. Sci. Rep. 6. doi: 10.1038/srep32073

Chen, Y., Zhao, X., Li, G., Kumar, S., Sun, Z., Li, Y., et al. (2021). Genome-wide identification of the nramp gene family in spirodela polyrhiza and expression analysis under cadmium stress. Int. J. Mol. Sci. 22. doi: 10.3390/ijms22126414

Clemens, S., and Ma, J. F. (2016). Toxic Heavy Metal and Metalloid Accumulation in Crop Plants and Foods. Annu. Rev. Plant Biol. 67, 489–512. doi: 10.1146/annurev-arplant-043015-112301

Curie, C., Alonso, J. M., Jean, M. LE, Ecker, J. R., and Briat, J.-F. (2000). Involvement of NRAMP1 from Arabidopsis thaliana in iron transport. Biochem. J. 347, 749–755. doi: 10.1042/bj3470749

Eastman, P., Swails, J., Chodera, J. D., McGibbon, R. T., Zhao, Y., Beauchamp, K. A., et al. (2017). OpenMM 7: Rapid development of high performance algorithms for molecular dynamics. PLoS Comput. Biol. 13. doi: 10.1371/journal.pcbi.1005659

Edgar, R. C. (2004). MUSCLE: Multiple sequence alignment with high accuracy and high throughput. Nucleic. Acids Res. 32, 1792–1797. doi: 10.1093/nar/gkh340

EFSA (2009). Cadmium in food - Scientific opinion of the Panel on Contaminants in the Food Chain. EFSA Journal 7. doi: 10.2903/j.efsa.2009.980

Ehrnstorfer, I. A., Geertsma, E. R., Pardon, E., Steyaert, J., and Dutzler, R. (2014). Crystal structure of a SLC11 (NRAMP) transporter reveals the basis for transition-metal ion transport. Nat. Struct. Mol. Biol. 21, 990–996. doi: 10.1038/nsmb.2904

Eisenberg, D., Weiss, R. M., and Terwilliger, T. C. (1984). The hydrophobic moment detects periodicity in protein hydrophobicity. Proc. Nadl. Acad. Sci. USA 81, 140–144. doi: 10.1073/pnas.81.1.140

Fu, D., Zhang, Z., Wallrad, L., Wang, Z., Höller D, S., Ju, C., et al. (2022). Ca ^2+^-dependent phosphorylation of NRAMP1 by CPK21 and CPK23 facilitates manganese uptake and homeostasis in Arabidopsis. doi: 10.1073/pnas.2204574119

Gao, H., Xie, W., Yang, C., Xu, J., Li, J., Wang, H., et al. (2018). NRAMP2, a trans-Golgi network-localized manganese transporter, is required for Arabidopsis root growth under manganese deficiency. New Phytologist 217, 179–193. doi: 10.1111/nph.14783

García-Esquinas, E., Pollan, M., Tellez-Plaza, M., Francesconi, K. A., Goessler, W., Guallar, E., et al. (2014). Cadmium exposure and cancer mortality in a prospective cohort: The strong heart study. Environ Health Perspect. 122, 363–370. doi: 10.1289/ehp.1306587

Gietz, R. D., and Schiestl, R. H. (2007). High-efficiency yeast transformation using the LiAc/SS carrier DNA/PEG method. Nat. Protoc. 2, 31–34. doi: 10.1038/nprot.2007.13

Gill, S. S., and Tuteja, N. (2011). Cadmium stress tolerance in crop plants: Probing the role of sulfur. Plant Signal Behav. 6, 215–222. doi: 10.4161/psb.6.2.14880

Guerra, J. V. da S., Ribeiro-Filho, H. V., Jara, G. E., Bortot, L. O., Pereira, J. G. de C., and Lopes-de-Oliveira, P. S. (2021). pyKVFinder: an efficient and integrable Python package for biomolecular cavity detection and characterization in data science. BMC Bioinformatics 22. doi: 10.1186/s12859-021-04519-4

Guerra, J. V. S., Ribeiro-Filho, H. V., Pereira, J. G. C., and Lopes-De-Oliveira, P. S. (2023). KVFinder-web: A web-based application for detecting and characterizing biomolecular cavities. Nucleic Acids Res. 51, W289–W297. doi: 10.1093/nar/gkad324

Guo, J., Long, L., Chen, A., Dong, X., Liu, Z., Chen, L., et al. (2022). Tonoplast-localized transporter ZmNRAMP2 confers root-to-shoot translocation of manganese in maize. Plant Physiol. 190, 2601– 2616. doi: 10.1093/plphys/kiac434

Hekkelman, M. L., de Vries, I., Joosten, R. P., and Perrakis, A. (2023). AlphaFill: enriching AlphaFold models with ligands and cofactors. Nat. Methods 20, 205–213. doi: 10.1038/s41592-022-01685-y

Honma, T., Shiratori, Y., Ohba, H., Tsuchida, T., Makino, T., Abe, T., et al. (2017). Concentrations of nutrient content in rice variety Koshihikari Kan No.1 and risk estimation of incidence of brown spot disease in different paddy fields. JPNSS 88, 213–220. doi: 10.20710/dojo.88.3_213

Hutchinson, T. C., Meema, K. M., Page, A. L., Chang, A. C., and El-Amamy, M. (1987). Lead, Mercury, Cadmium and Arsenic in the Environment., eds. T. C. Hutchinson and K. M. Meema. Published on behalf of the Scientific Committee on Problems of the Environment of the International Council of Scientific Unions by Wiley.

Ishikawa, S., Ishimaru, Y., Igura, M., Kuramata, M., Abe, T., Senoura, T., et al. (2012). Ion-beam irradiation, gene identification, and marker-assisted breeding in the development of low-cadmium rice. Proc. Natl. Acad. Sci. U S A 109, 19166–19171. doi: 10.1073/pnas.1211132109

Ishimaru, Y., Takahashi, R., Bashir, K., Shimo, H., Senoura, T., Sugimoto, K., et al. (2012). Characterizing the role of rice NRAMP5 in Manganese, Iron and Cadmium Transport. Sci. Rep. 2. doi: 10.1038/srep00286

Järup L., and Alfvén T. (2004). Low level cadmium exposure, renal and bone effects - the OSCAR study. BioMetals 17, 505–509.

Jones, D. T., Taylor, W. R., and Thornton, J. M. (1992). The rapid generation of mutation data matrices from protein sequences. Comput. Appl. Biosci. 8, 275–282. doi: 10.1093/bioinformatics/8.3.275

Julin, B., Wolk, A., Johansson, J. E., Andersson, S. O., Andrén, O., and Kesson, A. (2012). Dietary cadmium exposure and prostate cancer incidence: A population-based prospective cohort study. Br. J. Cancer 107, 895–900. doi: 10.1038/bjc.2012.311

Jumper, J., Evans, R., Pritzel, A., Green, T., Figurnov, M., Ronneberger, O., et al. (2021). Highly accurate protein structure prediction with AlphaFold. Nature 596, 583–589. doi: 10.1038/s41586-021-03819-2

Jurcik, A., Bednar, D., Byska, J., Marques, S. M., Furmanova, K., Daniel, L., et al. (2018). CAVER Analyst 2.0: Analysis and visualization of channels and tunnels in protein structures and molecular dynamics trajectories. Bioinformatics 34, 3586–3588. doi: 10.1093/bioinformatics/bty386

Kubier, A., Wilkin, R. T., and Pichler, T. (2019). Cadmium in soils and groundwater: A review. Applied Geochemistry 108. doi: 10.1016/j.apgeochem.2019.104388

Kuramata, M., Abe, T., Tanikawa, H., Sugimoto, K., and Ishikawa, S. (2022). A weak allele of OsNRAMP5 confers moderate cadmium uptake while avoiding manganese deficiency in rice. J. Exp. Bot. 73, 6475–6489. doi: 10.1093/jxb/erac302

Li, H., Wang, N., Hu, W., Yan, W., Jin, X., Yu, Y., et al. (2022). ZmNRAMP4 Enhances the Tolerance to Aluminum Stress in Arabidopsis thaliana. Int. J. Mol. Sci. 23. doi: 10.3390/ijms23158162

Li, J., Liu, Y., Kong, L., Xu, E., Zou, Y., Zhang, P., et al. (2023). An intracellular transporter OsNRAMP7 is required for distribution and accumulation of iron into rice grains. Plant Science 336. doi: 10.1016/j.plantsci.2023.111831

Li, J., Wang, L., Zheng, L., Wang, Y., Chen, X., and Zhang, W. (2018). A functional study identifying critical residues involving metal transport activity and selectivity in natural resistance-associated macrophage protein 3 in arabidopsis thaliana. Int. J. Mol. Sci. 19. doi: 10.3390/ijms19051430

Li, J., Wang, Y., Zheng, L., Li, Y., Zhou, X., Li, J., et al. (2019). The Intracellular Transporter AtNRAMP6 Is Involved in Fe Homeostasis in Arabidopsis. Front. Plant Sci. 10. doi: 10.3389/fpls.2019.01124

Lin, Q., Zong, Y., Xue, C., Wang, S., Jin, S., Zhu, Z., et al. (2020). Prime genome editing in rice and wheat. Nat. Biotechnol. 38, 582–585. doi: 10.1038/s41587-020-0455-x

Lu, M., Yang, G., Li, P., Wang, Z., Fu, S., Zhang, X., et al. (2018). Bioinformatic and functional analysis of a key determinant underlying the substrate selectivity of the al transporter, Nrat1. Front. Plant. Sci. 9. doi: 10.3389/fpls.2018.00606

Mirdita, M., Schütze, K., Moriwaki, Y., Heo, L., Ovchinnikov, S., and Steinegger, M. (2022). ColabFold: making protein folding accessible to all. Nat. Methods 19, 679–682. doi: 10.1038/s41592-022-01488-1

Mirdita, M., Steinegger, M., and Söding, J. (2019). MMseqs2 desktop and local web server app for fast, interactive sequence searches. Bioinformatics 35, 2856–2858. doi: 10.1093/bioinformatics/bty1057

Nazir, M. M., Noman, M., Ahmed, T., Ali, S., Ulhassan, Z., Zeng, F., et al. (2022). Exogenous calcium oxide nanoparticles alleviate cadmium toxicity by reducing Cd uptake and enhancing antioxidative capacity in barley seedlings. J. Hazard Mater. 438. doi: 10.1016/j.jhazmat.2022.129498

Nogawa, K., and Kido, T. (1993). Envrolmnental Health Biological monitoring of cadmium exposure in itai-itai disease epidemiology. Int. Arch. Occup. Environ. Health 65, 543–546. doi: 10.1007/BF00381306

Peris-Peris, C., Serra-Cardona, A., Śanchez-Sanuy, F., Campo, S., Ariño, J., and Segundo, B. S. (2017). Two NRAMP6 isoforms function as iron and manganese transporters and contribute to disease resistance in rice. Molecular Plant-Microbe Interactions 30, 385–398. doi: 10.1094/MPMI-01-17-0005-R

Podar, D., Scherer, J., Noordally, Z., Herzyk, P., Nies, D., and Sanders, D. (2012). Metal selectivity determinants in a family of transition metal transporters. J. Biol. Chem. 287, 3185–3196. doi: 10.1074/jbc.M111.305649

Pottier, M., Oomen, R., Picco, C., Giraudat, J., Scholz-Starke, J., Richaud, P., et al. (2015). Identification of mutations allowing Natural Resistance Associated Macrophage Proteins (NRAMP) to discriminate against cadmium. Plant Journal 83, 625–637. doi: 10.1111/tpj.12914

Qing, Y., Li, Y., Cai, X., He, W., Liu, S., Ji, Y., et al. (2023). Assessment of Cadmium Concentrations in Foodstuffs and Dietary Exposure Risk Across China: A Metadata Analysis. Expo. Health 15, 951– 961. doi: 10.1007/s12403-022-00530-z

Qu, Z., and Nakanishi, H. (2023). Amino acid residues of the metal transporter OsNRAMP5 responsible for cadmium absorption in rice. Plants 12. doi: 10.3390/plants12244182

Ray, S., Berry, S. P., Wilson, E. A., Zhang, C. H., Shekhar, M., Singharoy, A., et al. (2023). High-resolution structures with bound Mn^2+^ and Cd^2+^ map the metal import pathway in an Nramp transporter. Elife 12. doi: 10.7554/eLife.84006

Rentsch, D., Laloi, M., Rouhara, I., Schmelzer, E., Delrot, S., and Frommer, W. B. (1995). NTR1 encodes a high affinity oligopeptide transporter in Arabidopsis. FEBS Lett 370, 264–268. doi: 10.1016/0014-5793(95)00853-2

Ritz, C., Baty, F., Streibig, J. C., and Gerhard, D. (2015). Dose-response analysis using R. PLoS One 10. doi: 10.1371/journal.pone.0146021

Sasaki, A., Yamaji, N., Yokosho, K., and Ma, J. F. (2012). Nramp5 is a major transporter responsible for manganese and cadmium uptake in rice. Plant Cell 24, 2155–2167. doi: 10.1105/tpc.112.096925

Socha, A. L., and Guerinot, M. Lou (2014). Mn-euvering manganese: The role of transporter gene family members in manganese uptake and mobilization in plants. Front. Plant Sci. 5. doi: 10.3389/fpls.2014.00106

Staessen, J. A., Roels, H. A., Emelianov, D., Kuznetsova, T., Thijs, L., Vangronsveld, J., et al. (1999). Environmental exposure to cadmium, forearm bone density, and risk of fractures: prospective population study. THE LANCET 353, 1140–44. doi: 10.1016/s0140-6736(98)09356-8.

Sui, F. Q., Chang, J. D., Tang, Z., Liu, W. J., Huang, X. Y., and Zhao, F. J. (2018). Nramp5 expression and functionality likely explain higher cadmium uptake in rice than in wheat and maize. Plant Soil 433, 377–389. doi: 10.1007/s11104-018-3849-5

Sun, G. L., Reynolds, E. E., and Belcher, A. M. (2019). Designing yeast as plant-like hyperaccumulators for heavy metals. Nat. Commun. 10. doi: 10.1038/s41467-019-13093-6

Takahashi, R., Ishimaru, Y., Senoura, T., Shimo, H., Ishikawa, S., Arao, T., et al. (2011). The OsNRAMP1 iron transporter is involved in Cd accumulation in rice. J. Exp. Bot. 62, 4843–4850. doi: 10.1093/jxb/err136

Tamura, K., Stecher, G., and Kumar, S. (2021). MEGA11: Molecular Evolutionary Genetics Analysis Version 11. Mol. Biol. Evol. 38, 3022–3027. doi: 10.1093/molbev/msab120

Tao, R., Wang, Y., Jiao, Y., Hu, Y., Li, L., Jiang, L., et al. (2022). Bi-PE: bi-directional priming improves CRISPR/Cas9 prime editing in mammalian cells. Nucleic Acids Res. 50, 6423–6434. doi: 10.1093/nar/gkac506

Thomine, S., Wang, R., Ward, J. M., Crawford, N. M., and Schroeder, J. I. (2000). Cadmium and iron transport by members of a plant metal transporter family in Arabidopsis with homology to Nramp genes. Proc. Natl. Acad. Sci. USA 97, 4991–4996. doi: 10.1073/pnas.97.9.4991

Tim, N., Michelle, P., Janneke, H., Harry, A. R., Hilde, C., Lutgarde, T., et al. (2006). Environmental exposure to cadmium and risk of cancer: a prospective population-based study. Lancet Oncol. 7, 119–126. doi: 10.1016/S1470-2045(06)70545-9

Tiwari, M., Sharma, D., Dwivedi, S., Singh, M., Tripathi, R. D., and Trivedi, P. K. (2014). Expression in Arabidopsis and cellular localization reveal involvement of rice NRAMP, OsNRAMP1, in arsenic transport and tolerance. Plant Cell Environ. 37, 140–152. doi: 10.1111/pce.12138

Vervaet, B. A., D’Haese, P. C., and Verhulst, A. (2017). Environmental toxin-induced acute kidney injury. Clin Kidney J 10, 747–758. doi: 10.1093/ckj/sfx062

Wallin, M., Barregard, L., Sallsten, G., Lundh, T., Sundh, D., Lorentzon, M., et al. (2021). Low-level cadmium exposure is associated with decreased cortical thickness, cortical area and trabecular bone volume fraction in elderly men: The MrOS Sweden study. Bone 143. doi: 10.1016/j.bone.2020.115768

World Health Organization (2006). Evaluation of Certain Food Contaminants : Sixty-fourth Report of the Joint FAO/WHO Expert Committee on Food Additives. Geneva: World Health Organization.

World Health Organization (2011). Evaluation of certain food additives and contaminants : seventy-third report of the Joint FAO/WHO Expert Committee on Food Additives. Geneva, Switzerland: World Health Organization.

World Health Organization (2022). Guidelines for drinking-water quality: Fourth edition incorporating the first and second addenda., ed. World Health Organization.

Xia, J., Yamaji, N., Kasai, T., and Ma, J. F. (2010). Plasma membrane-localized transporter for aluminum in rice. Proc. Natl. Acad. Sci. U S A 107, 18381–18385. doi: 10.1073/pnas.1004949107

Yamaji, N., Sasaki, A., Xia, J. X., Yokosho, K., and Ma, J. F. (2013). A node-based switch for preferential distribution of manganese in rice. Nat. Commun. 4. doi: 10.1038/ncomms3442

Yang, M., Zhang, W., Dong, H., Zhang, Y., Lv, K., Wang, D., et al. (2013). OsNRAMP3 is a vascular bundles-specific manganese transporter that is responsible for manganese distribution in rice. PLoS One 8. doi: 10.1371/journal.pone.0083990

Yang, M., Zhang, Y., Zhang, L., Hu, J., Zhang, X., Lu, K., et al. (2014). OsNRAMP5 contributes to manganese translocation and distribution in rice shoots. J. Exp. Bot. 65, 4849–4861. doi: 10.1093/jxb/eru259

Yohannan, S., Yang, D., Faham, S., Boulting, G., Whitelegge, J., and Bowie, J. U. (2004). Proline substitutions are not easily accommodated in a membrane protein. J. Mol. Biol. 341, 1–6. doi: 10.1016/j.jmb.2004.06.025

Zhang, X., Chen, D., Zhong, T., Zhang, X., Cheng, M., and Li, X. (2015). Assessment of cadmium (Cd) concentration in arable soil in China. Environ. Sci. Pollut. Res. 22, 4932–4941. doi: 10.1007/s11356-014-3892-6

Zhao, J., Yang, W., Zhang, S., Yang, T., Liu, Q., Dong, J., et al. (2018). Genome-wide association study and candidate gene analysis of rice cadmium accumulation in grain in a diverse rice collection. Rice 11. doi: 10.1186/s12284-018-0254-x

Zong, Y., Liu, Y., Xue, C., Li, B., Li, X., Wang, Y., et al. (2022). An engineered prime editor with enhanced editing efficiency in plants. Nat. Biotechnol. doi: 10.1038/s41587-022-01254-w

