## Supplementary Materials for "Engineering rice Nramp5 modifies cadmium and manganese uptake selectivity using yeast assay system"

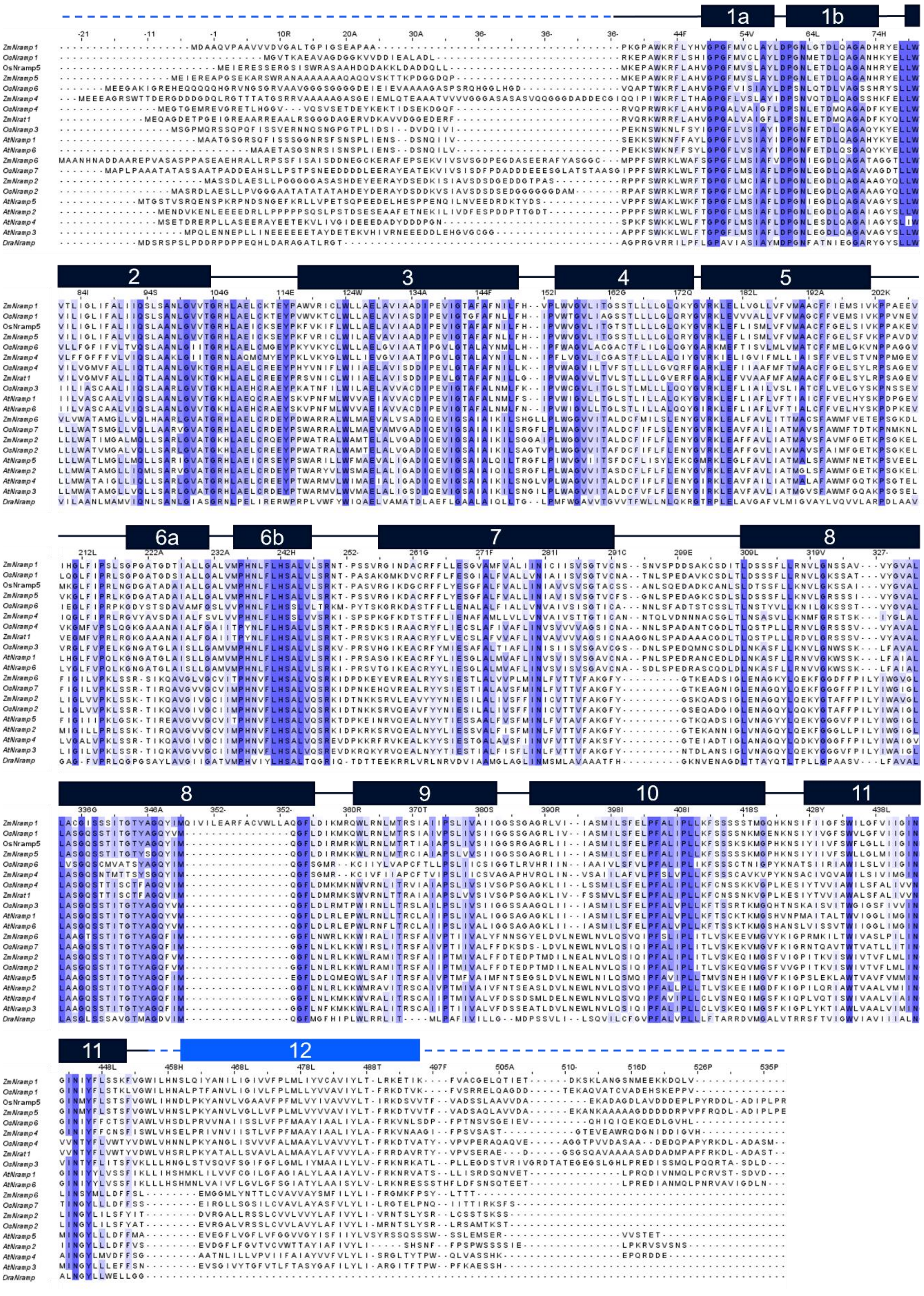

**Supplementary Figure 1. Amino acid sequences of Nramp proteins.** Amino acid sequences of 20 Nramps shown with numbers corresponding to OsNramp5. The amino acid conservation is shown in blue. The transmembrane region is schematically represented.

A

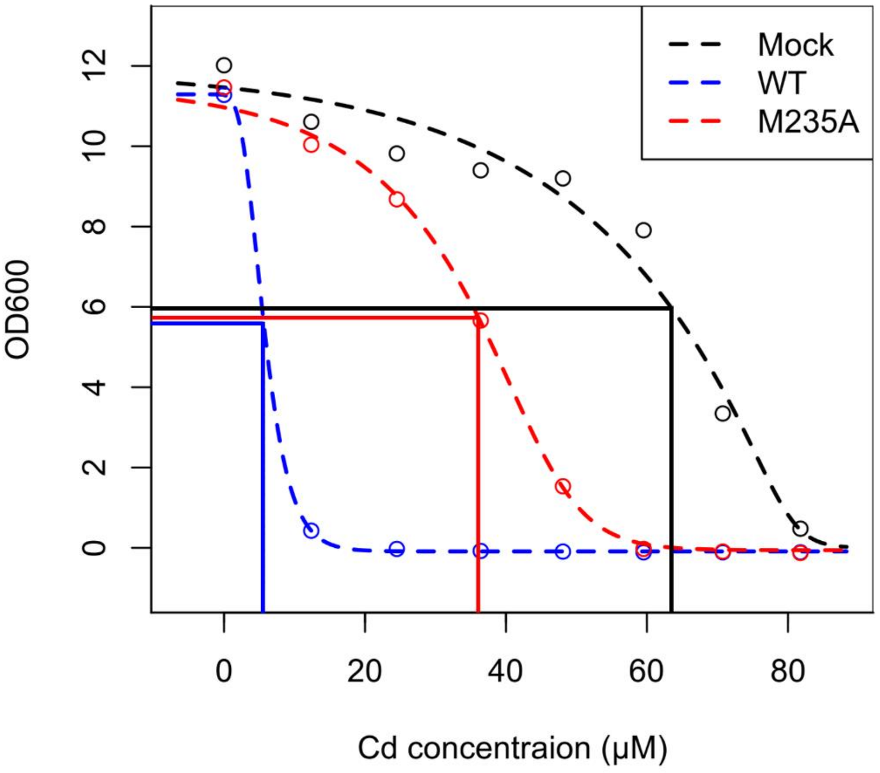


B

| | $IC_{50}$ | $1 / IC_{50}$ | Relative Cd uptake (%) |
|---|---|---|---|
| Mock | 63.5 | 0.016 | 0.0 |
| WT | 5.6 | 0.178 | 100.0 |
| M235A | 36.1 | 0.028 | 7.4 |

**Supplementary Figure 2. Example of estimation of the metal uptake efficiency. (A)** Cd-sensitive strain (*Δycf1*) containing the plasmid (WT, M235A, or EV) was incubated in various Cd concentrations (0, 20, 40, 60, or 80 μM). The growth was measured by the OD600 and the maximum likelihood curves are depicted to estimate the $IC_{50}$. **(B)** Metal uptake efficiency estimated as the inversed score of $IC_{50}$ ($1/ IC_{50}$ ) with adjustment of the score of WT and EV as 100 and 0, respectively.

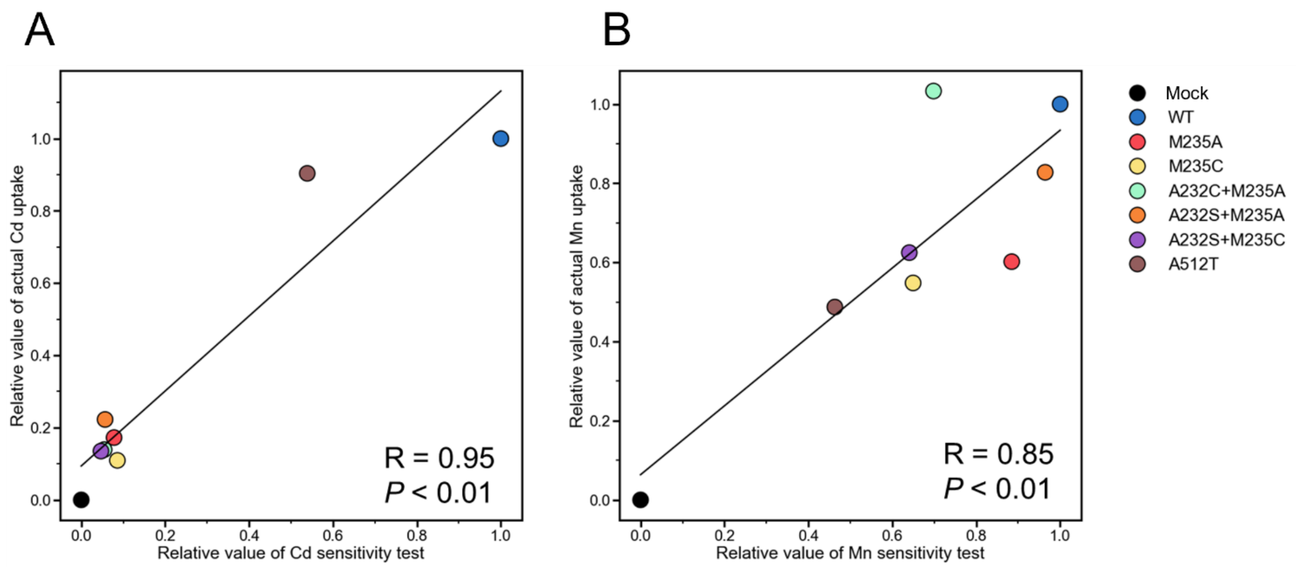


**Supplementary Figure 3. Compatibilities between the result of the yeast assay system and ICP-OES. (A)** Relative Cd uptake value obtained by the yeast assay system (Table 2) and scatter plot of the direct measurement using ICP-OES (Table 3). Pearson's correlation coefficient R and the *P* values are shown to estimate the compatibility. **(B)** Relative Mn uptake values obtained from the two methodologies in (A).

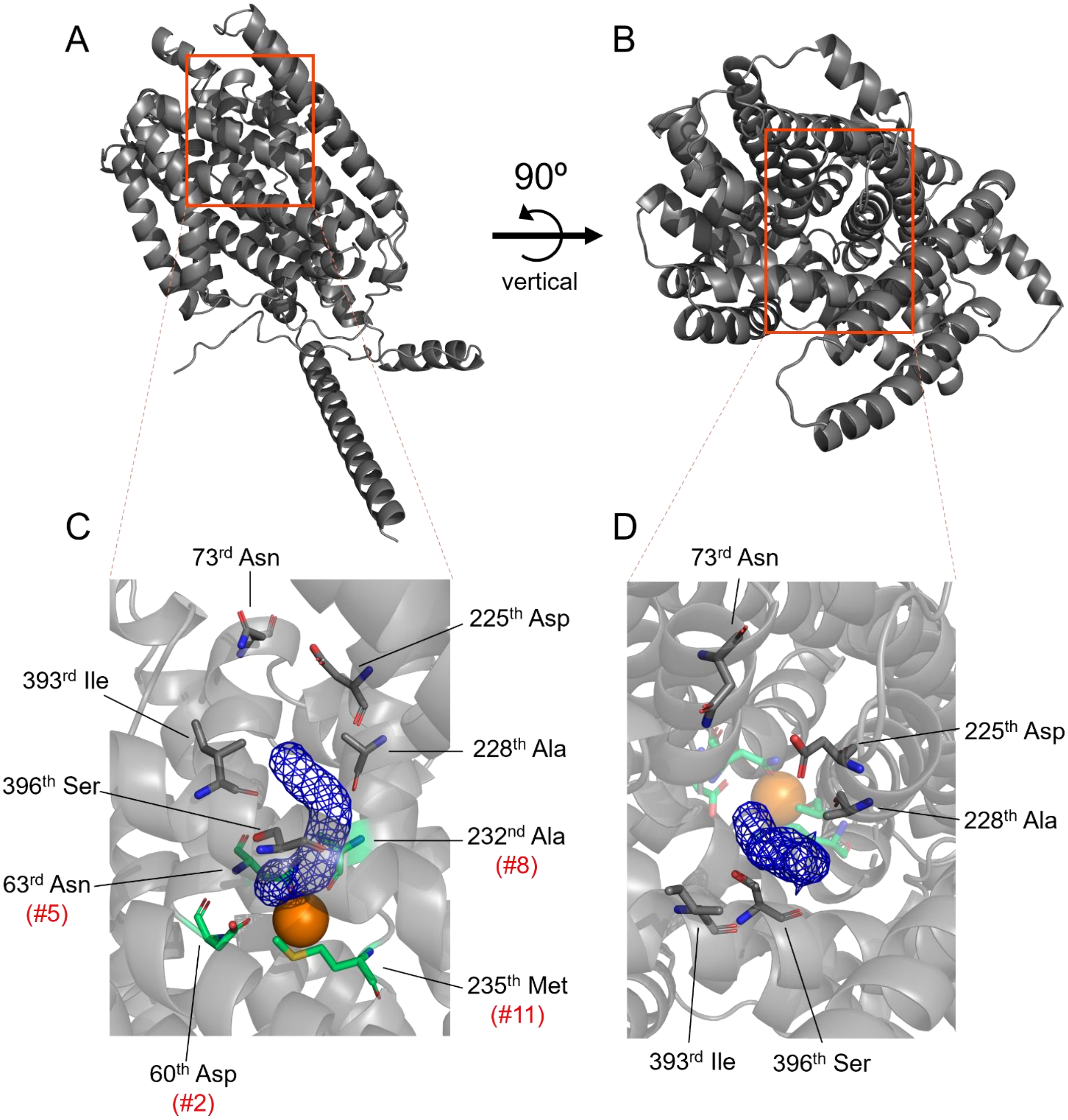


**Supplementary Figure 4. Predicted Mn transport pathway of OsNramp5.** Predicted structure of OsNramp5 viewed from the **(A)** membrane side and **(B)** extracellular metal entry side. **(C)** Predicted transport tunnel from the membrane side. The transport tunnel and $Mn^{2+}$ ions are shown by blue mesh and orange spheres, respectively. The key residues for the metal uptake efficiency (#2, #5, #8, and #11) are shown by green sticks, together with the amino acid residues that have been demonstrated to be involved in the metal uptake efficiency and selectivity of other Nramps. **(D)** Predicted transport tunnel from the extracellular or metal entry side shown as in (C).

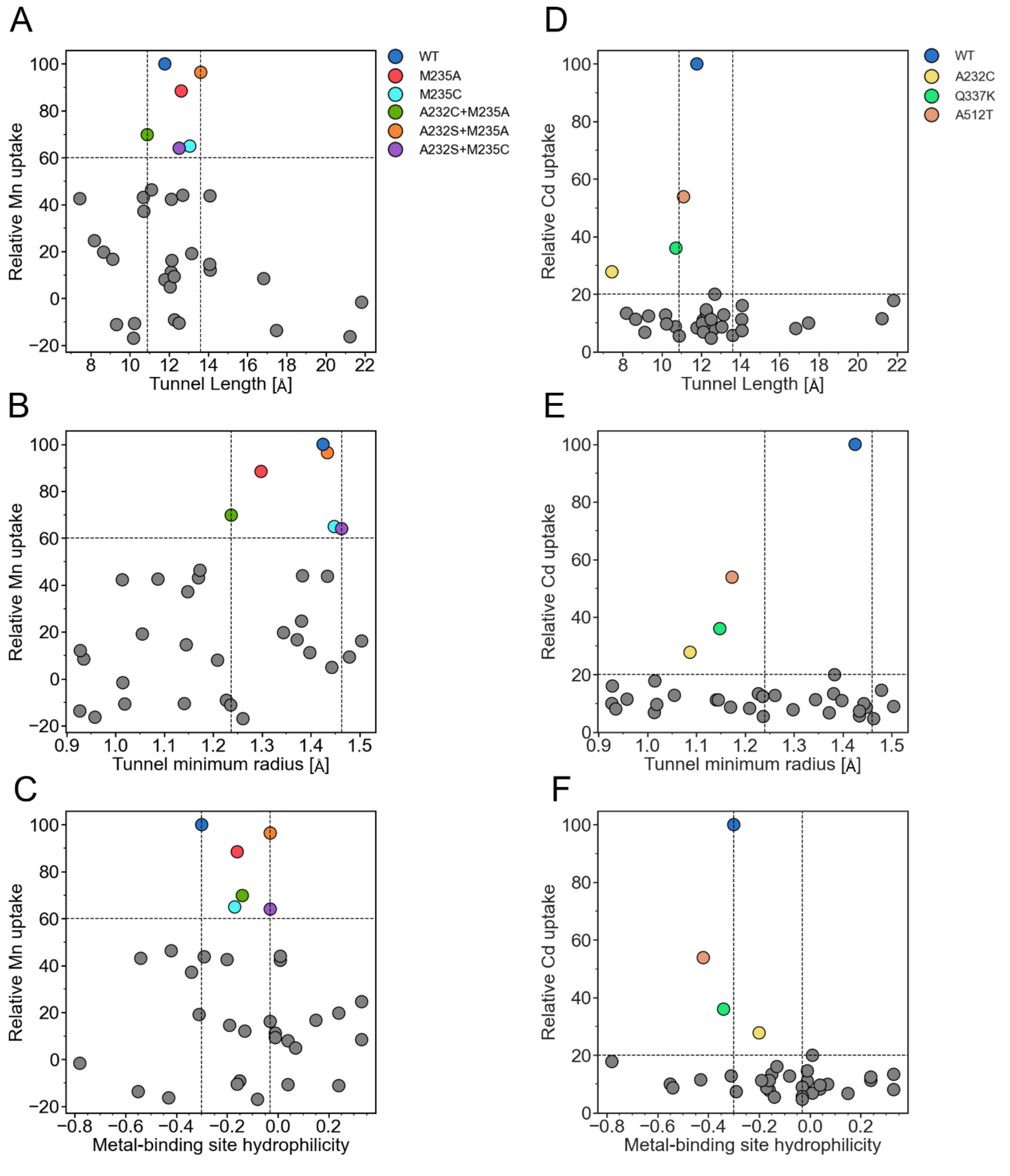


**Supplementary Figure 5. Relationship between the relative metal uptake efficiency and spatial/chemical properties of the predicted 3D structure. (A)** Relative Mn uptake of the OsNramp5 and 33mutants in this study (Table 1 and 2) scatter-plotted according to the tunnel length of the predicted structure. The proteins that showed more than 60% Mn uptake efficiency were positioned in the tunnel length between 10.88 and 13.61 Å. Scatter plots of relative Mn uptake and (B) tunnel minimum radius and (C) metal-binding site hydrophilicity are shown, as well as for relative Cd uptake (D, E, and F).

**Supplementary Table 1.** Nramp genes for the phylogenetic analysis

| Gene | Accession | Locus |
|---|---|---|
| AtNramp1 | NP_178198 | AT1G80830 |
| AtNramp2 | NP_175157 | AT1G47240 |
| AtNramp3 | NP_179896 | AT2G23150 |
| AtNramp4 | NP_201534 | AT5G67330 |
| AtNramp5 | NP_193614 | AT4G18790 |
| AtNramp6 | NP_173048 | AT1G15960 |
| OsNramp1 | NP_001389971 | Os07g0258400 |
| OsNramp2 | NP_001404287 | Os03g0208500 |
| OsNramp3 | NP_001411161 | Os06g0676000 |
| OsNramp4 | NP_001396197 | Os02g0131800 |
| OsNramp5 | NP_001389970 | Os07g0257200 |
| OsNramp6 | NP_001393044 | Os01g0503400 |
| OsNramp7 | NP_001410524 | Os12g0581600 |
| ZmNramp1 | XP_008670084 | GRMZM2G366919 |
| ZmNramp2 | NP_001150280 | GRMZM2G178190 |
| ZmNramp4 | XP_008670762 | GRMZM2G322844 |
| ZmNramp5 | NP_001354866 | GRMZM2G147560 |
| ZmNramp6 | XP_008665146 | GRMZM2G025680 |
| ZmNrat1 | NP_001334019 | GRMZM2G168747 |
| ScaNramp | WP_002452964 | - |
| DraNramp | Q9RTP8 | - |

**Supplementary Table 2.** Sequence of primers

| Primer name | Forward(5'-3') | Reverse(3'-5') |
|---|---|---|
| OsNramp5_GA | AAAAATATACCCCAGCCATGGAGATTGAGAGAGAGAGC | GAAGAAGTCCAAAGCTGACACCCTTGTCGATCGATC |
| M235A | CTTGTCGCGCCCCACAATCTGTTCTTG | GTGGGGCGCGACAAGAGCTCCGAGGAG |
| M235C | CTTGTCTGCCCCCACAATCTGTTCTTG | GTGGGGGCAGACAAGAGCTCCGAGGAG |
| M235D | CTTGTCGACCCCCACAATCTGTTCTTG | GTGGGGGTCGACAAGAGCTCCGAGGAG |
| M235E | CTTGTCGAGCCCCACAATCTGTTCTTG | GTGGGGCTCGACAAGAGCTCCGAGGAG |
| M235F | CTTGTCTTCCCCCACAATCTGTTCTTG | GTGGGGGAAGACAAGAGCTCCGAGGAG |
| M235G | CTTGTCGGGCCCCACAATCTGTTCTTG | GTGGGGCCCGACAAGAGCTCCGAGGAG |
| M235H | CTTGTCCATCCCCACAATCTGTTCTTG | GTGGGGATGGACAAGAGCTCCGAGGAG |
| M235I | CTTGTCATACCCCACAATCTGTTCTTG | GTGGGGTATGACAAGAGCTCCGAGGAG |
| M235K | CTTGTCAAGCCCCACAATCTGTTCTTG | GTGGGGCTTGACAAGAGCTCCGAGGAG |
| M235L | CTTGTCCTGCCCCACAATCTGTTCTTG | GTGGGGCAGGACAAGAGCTCCGAGGAG |
| M235N | CTTGTCAATCCCCACAATCTGTTCTTG | GTGGGGATTGACAAGAGCTCCGAGGAG |
| M235P | CTTGTCCCGCCCCACAATCTGTTCTTG | GTGGGGCGGGACAAGAGCTCCGAGGAG |
| M235Q | CTTGTCCAGCCCCACAATCTGTTCTTG | GTGGGGCTGGACAAGAGCTCCGAGGAG |
| M235R | CTTGTCAGGCCCCACAATCTGTTCTTG | GTGGGGCCTGACAAGAGCTCCGAGGAG |
| M235S | CTTGTCAGTCCCCACAATCTGTTCTTG | GTGGGGACTGACAAGAGCTCCGAGGAG |
| M235T | CTTGTCACGCCCCACAATCTGTTCTTG | GTGGGGCGTGACAAGAGCTCCGAGGAG |
| M235V | CTTGTCGTGCCCCACAATCTGTTCTTG | GTGGGGCACGACAAGAGCTCCGAGGAG |
| M235W | CTTGTCTGGCCCCACAATCTGTTCTTG | GTGGGGCCAGACAAGAGCTCCGAGGAG |
| M235Y | CTTGTCTACCCCCACAATCTGTTCTTG | GTGGGGGTAGACAAGAGCTCCGAGGAG |
| A232S | CTCGGAAGTCTTGTCATGCCCCACAATCTGTTC | GACAAGACTTCCGAGGAGGGCAATGGCGTCGGC |
| A232V | CTCGGAGTGCTTGTCATGCCCCACAATCTGTTC | GACAAGCACTCCGAGGAGGGCAATGGCGTCGGC |
| A232C | CTCGGATGCCTTGTCATGCCCCACAATCTGTTC | GACAAGGCATCCGAGGAGGGCAATGGCGTCGGC |
| A232M | CTCGGAATGCTTGTCATGCCCCACAATCTGTTC | GACAAGCATTCCGAGGAGGGCAATGGCGTCGGC |
| A232S+M235A | GGAAGTCTTGTCGCGCCCCACAATCTGTTC | GGGCGCGACAAGACTTCCGAGGAGGGCAAT |
| A232V+M235A | GGAGTGCTTGTCGCGCCCCACAATCTGTTC | GGGCGCGACAAGCACTCCGAGGAGGGCAAT |
| A232C+M235A | GGATGCCTTGTCGCGCCCCACAATCTGTTC | GGGCGCGACAAGGCATCCGAGGAGGGCAAT |
| A232M+M235A | GGAATGCTTGTCGCGCCCCACAATCTGTTC | GGGCGCGACAAGCATTCCGAGGAGGGCAAT |
| A232S+M235C | GGAAGTCTTGTCTGCCCCCACAATCTGTTC | GGGGCAGACAAGACTTCCGAGGAGGGCAAT |
| A232V+M235C | GGAGTGCTTGTCTGCCCCCACAATCTGTTC | GGGGCAGACAAGCACTCCGAGGAGGGCAAT |
| A232C+M235C | GGATGCCTTGTCTGCCCCCACAATCTGTTC | GGGGCAGACAAGGCATCCGAGGAGGGCAAT |
| A232M+M235C | GGAATGCTTGTCTGCCCCCACAATCTGTTC | GGGGCAGACAAGCATTCCGAGGAGGGCAAT |
| Q337K | TCTGGGAAGAGCTCCACTATTACCGGC | GGAGCTCTTCCCAGATGCCAACAGTGC |
| A512T | GAGAAGACCGACGCCGGCGACCTCGCC | GGCGTCGGTCTTCTCGGCGTCGACGAC |

**Supplementary Table 3.** Properties of predicted transport pathways and metal-binding site

| Plasmid | Tunnel length [Å] | Tunnel minimum radius [Å] | Hydropathy around metal-binding site |
|---|---|---|---|
| WT | 11.78 | 1.43 | -0.30 |
| Q337K | 10.71 | 1.15 | -0.34 |
| A512T | 11.10 | 1.17 | -0.42 |
| M235A | 12.62 | 1.30 | -0.16 |
| M235C | 13.05 | 1.45 | -0.17 |
| M235D | 8.19 | 1.38 | 0.33 |
| M235E | 12.10 | 1.40 | -0.01 |
| M235F | 17.48 | 0.93 | -0.55 |
| M235G | 11.79 | 1.21 | 0.04 |
| M235H | 8.66 | 1.34 | 0.24 |
| M235I | 12.27 | 1.23 | -0.15 |
| M235K | 16.83 | 0.94 | 0.33 |
| M235L | 12.14 | 1.50 | -0.03 |
| M235N | 9.13 | 1.37 | 0.15 |
| M235P | 10.18 | 1.26 | -0.08 |
| M235Q | 9.32 | 1.24 | 0.24 |
| M235R | - | - | - |
| M235S | 12.06 | 1.44 | 0.07 |
| M235T | 12.26 | 1.48 | -0.01 |
| M235V | 12.51 | 1.14 | -0.16 |
| M235W | 21.82 | 1.02 | -0.78 |
| M235Y | 21.23 | 0.96 | -0.43 |
| A232C | 7.44 | 1.09 | -0.20 |
| A232M | 12.12 | 1.01 | 0.01 |
| A232S | 12.70 | 1.38 | 0.01 |
| A232V | 10.69 | 1.17 | -0.54 |
| A232C+M235A | 10.88 | 1.24 | -0.14 |
| A232M+M235A | 14.10 | 0.93 | -0.13 |
| A232S+M235A | 13.61 | 1.43 | -0.03 |
| A232V+M235A | 14.07 | 1.15 | -0.19 |
| A232C+M235C | 14.08 | 1.43 | -0.29 |
| A232M+M235C | 10.24 | 1.02 | 0.04 |
| A232S+M235C | 12.51 | 1.46 | -0.03 |
| A232V+M235C | 13.15 | 1.06 | -0.31 |
